# Caliditerrarchaeota, a new sister to Nanohaloarchaeota, provides insights into the evolution of DPANN halophily

**DOI:** 10.1101/2025.08.05.668663

**Authors:** Joshua N. Hamm, Nina Dombrowski, Luis E. Valentin-Alvarado, Chris Greening, Tom A. Williams, Anja Spang

## Abstract

The Nanohaloarchaeota are a clade of halophilic symbionts with small cells and genomes. Originally placed within the Euryarchaeota, they are now widely thought to belong to the DPANN archaea. However, the evolution of this clade and its phylogenetic placement within DPANN remain poorly understood.

We applied phylogenetic and comparative genomic analyses to assess the evolutionary relationship and genome evolution of the Nanohaloarchaeota and related DPANN lineages. Our phylogenetic analyses resolve Nanohaloarchaeota as a sister group to a phylum-level lineage referred to as EX4484-52, together forming a clade with Aenigmatarchaeota. Representatives of EX4484-52, for which we propose the name Caliditerrarchaeota, have an anaerobic, thermophilic lifestyle but do not appear to be adapted to high salt concentrations.

Using gene tree-species tree reconciliations, we investigated the origin of halophily across these archaeal lineages revealing that adaptations to high-salt appear to have evolved on the branch to Nanohaloarchaeota after their divergence from Aenigmatarchaeota and Caliditerrarchaeota. In agreement with recent work, we also identify hallmarks of halophily in another order-level lineage within the Aenigmatarchaeota (Haloaenigmatarchaeaceae) which appears to represent a second independent adaptation of a DPANN clade to halophily. The two halophilic DPANN lineages are inferred to have distinct sets of proteins that enable them to live in environments with high salt levels. Notably, phylogenetic analyses reveal a dominant signal of gene transfers between Haloaenigmatarchaeaceae and Halarchaeoplasmatales, indicating a potential host-symbiont relationship.

This work provides the first detailed investigation of the enigmatic Caliditerrarchaeota, and new insights into the evolution of halophilic lifestyles within DPANN.

## Introduction

The DPANN archaea, named after the first representatives of this group [1–5], comprise at least 10 phylum-level lineages (out of 21 archaeal phyla in total) [6]. Yet only 3 DPANN phyla have cultivated representatives, i.e. the Nanoarchaeota [7–10], Nanohaloarchaeota [11–13] and Micrarchaeota [14,15]. Currently, the lifestyles of most groups are poorly understood. It has recently been suggested that the DPANN form at least two distinct major groups: Cluster 1, which includes the Altiarchaeota, Iainarchaeota, and Micrarchaeota and Cluster 2, including Nanohaloarchaeota, Nanoarchaeota, Aenigmatarchaeota, EX4484-52, SpSt-1190, and Undinarchaeota [4,6,16]. Distributed across the biosphere, Cluster 2 DPANN are generally characterised by reduced genomes and small cells, and are predicted to rely upon host organisms [3]. For instance, cultivated representatives of the Nanoarchaeota [7–10] and Nanohaloarchaeota [11–13] depend on archaeal host cells - i.e. members of the *Sulfolobales* and *Halobacteriales*, respectively - for growth and survival.

Until recently, the Nanohaloarchaeota were the only known DPANN lineage adapted to extreme salt conditions [17]. However, the recently-discovered Haloaenigmatarchaeaceae (within phylum Aenigmatarchaeota) are a second DPANN lineage that seems to have independently evolved a halophilic lifestyle [18]. Similar to other halophilic archaea, the Nanohaloarchaeota and Haloaenigmatarchaeaceae appear to employ a salt-in strategy in which high intracellular salt concentrations are tolerated via greater use of acidic amino acids in proteins, leading to a shift in proteome-wide amino acid composition [18–23]. Of particular importance are transporters involved in uptake of K^+^ (Such as the Kef-type transporter), and exporters of Na^+^ (such as the Na^+^/H^+^ antiporter) which maintain osmotic balance through selective import of K^+^ and export of Na^+^ [24]. Since its discovery, the phylogenetic placement of the Nanohaloarchaeota phylum has been a subject of debate. Initial data supported a sister relationship with the *Halobacteriales* [20] or Methanocellales [25], others recovered a sisterhood with other DPANN [5]. An increasing amount of work based on concatenated single-copy marker gene phylogenies implementing more realistic models of evolution has provided growing evidence that Nanohaloarchaeota indeed belong to the DPANN archaea [2,4,22,26–28]. The monophyletic clustering of nanohaloarchaeal and haloarchaeal proteins in single protein phylogenies is likely a result of protein compositional biases as a response to high salt-adaptation and increased rates of horizontal gene transfer (HGT) between these halophilic symbionts and their hosts [4,22,29].

Despite a better understanding of the phylogenetic placement of the Nanohaloarchaeota, a major limitation in assessing the evolution of halophily in Nanohaloarchaeota was the absence of closely related sister lineages with which to compare lifestyles and genetic content. Recently, the *Asbonarchaeaceae* were reported to be a deep branching, halophilic sister lineage to the *Nanosalinaceae* within the Nanohaloarchaeota phylum found in polyextreme environments [22]. Yet, the origin and nature of adaptations to a halophilic lifestyle in the Nanohaloarchaeota and their relationship with non-halophilic DPANN cluster 2 lineages, especially the Aenigmatarchaeota [5,30] and EX4484-52 phylum [31–33] remains to be elucidated. Gene tree-species tree reconciliation methods for ancestral genome reconstruction have been applied to the Nanohaloarchaeota previously but have focused on internal nodes within the nanohaloarchaeal lineage, leaving the transition from non-halophile to halophile unexplored [34,35]. Here, we used phylogenetics, comparative genomics, and gene trees-species tree reconciliation [36,37] to place the Nanohaloarchaeota within DPANN cluster 2 archaea, characterise their closest sister lineage EX4484-52, for which we propose the name Caliditerrarchaeota, and reconstruct the genome evolution of DPANN cluster 2 lineages to assess genomic signatures associated with the adaptation to a halophilic lifestyle. Our work identifies the Caliditerrarchaeota as primarily non-halophilic terrestrial thermophiles, and highlights DNA repair and potassium transport as likely key to halophily in DPANN. Increased rates of horizontal gene transfer between Haloaenigmatarchaeaceae and the Halarchaeoplasmatales point to a putative symbiont-host relationship between members of these clades.

## Methods

### Generation of dataset for phylogenetic and genomic analyses

To investigate the placement of halophilic lineages a previously established representative set of 607 archaeal genomes was used as a backbone [4]. To this, 44 publicly available genomes associated with the Nanohaloarchaeota-Aenigmatarchaeota clade, Hikarchaeota, and Thermoplasmatales were added (Supplementary Table 1) resulting in a final dataset of 651 genomes.

### Selection of phylogenetic marker genes for species tree reconstruction

To generate a reliable species tree and accurately place ACN lineages for downstream analyses, we applied a phylogenetic approach similar to Dombrowski et. al. 2020 [4] to identify suitable marker genes from a set of 151 possible markers (Supplementary Table 12). HMM profiles specific to the marker genes were used to identify homologs in the 651 genomes using a custom script Selection_of_Phylogenetic_Markers/Selecting_best_markers_151set.sh [38]) that incorporates the hmmsearch [39] algorithm from the HMMER v3.1b2 package. To reduce the possibility of distant paralogs being extracted during this process the set of HMM profiles used as query was expanded to include the entire TIGRFAM[40] database (4,528 profiles total). Subsequently, homologs to each of the 151 marker genes were parsed from the output based on best hit evaluated by e-value and bit-score and extracted from their respective genomes. Single gene alignments were performed for all 151 marker genes using MAFFT L-INS-i v7.407 [41] (settings:reorder) and trimmed using BMGE-1.12 [42] (settings:-tAA -m BLOSUM30 -h 0.55). Phylogenies were inferred for all genes using IQ-tree v2.1.2 [43] (settings: -m LG+G -bb 1000 -wbtl -bnni). Single gene phylogenies were assessed using a custom python script to rank markers based on their capacity to resolve established monophyletic taxa as previously described [4]. Reliability of markers was scored according to the number of total splits (total number of taxa placing outside the expected taxonomic member clade) as well as the number of splits normalised to species count within a specific clade. Marker genes in which Archaea were previously found to not be monophyletic [4] were removed. Split counts were used to rank the marker genes and concatenated alignments of the 25% and 50% top and lowest ranked genes were generated (Supplementary Table 12).

### Species tree reconstruction

Sequences on long branches were flagged and removed using a custom python script Selection_of_Phylogenetic_Markers/cut_gene_tree_v2.py [38] applied to trees generated for marker gene reliability assessment. Following long branch removal, remaining sequences were re-aligned and trimmed using MAFFT L-INS-i v7.407 [41] and BMGE-1.12 [42] as described above. Marker gene sets were concatenated using catfasta2phyml.pl (https://github.com/nylander/catfasta2phyml). To assess placement of halophilic lineages as well as the putative sister clade of the Nanohaloarchaeota (i.e. the Caliditerrarchaeota) multiple concatenated trees were generated:

1. Phylogenies using marker gene subsets: All four subsets of marker genes (top 25 and 50% and lowest 25 and 50%). Alignments were concatenated using catfasta2phyml.pl (https://github.com/nylander/catfasta2phyml) and trees generated using IQ-tree [43] (v2.1.2, settings: -m LG+C60+F+R -nt AUTO -bb 1000-alrt 1000).
2. Removal of compositionally heterogeneous sites: Concatenated alignments of the top 25 and 50% best marker genes were aligned with MAFFT L-INS-i v7.407 [41] and trimmed with BMGE-1.12 [42] as above. Alignments were then subjected to compositional site filtering using a custom perl script (https://github.com/novigit/davinciCode/blob/master/perl/alignment_pruner.pl) [44] in a stepwise manner to remove the 10, 30, and 50% most compositionally heterogeneous sites. Trees were reconstructed for all three conditions of the two marker subsets using IQ-tree [43] (v2.1.2, settings: -m LG+C60+F+R -nt AUTO - bb 1000 -alrt 1000).
3. Removal of fast-evolving sites: SlowFaster v1 [45] was used to remove fast evolving sites from the top 25 and 50% best marker gene alignments. Sites were filtered in a stepwise manner to remove the 10, 30, and 50% fastest evolving sites. Trees were reconstructed for all three conditions of the two marker subsets using IQ-tree [43] (v2.1.2, settings: -m LG+C60+F+R -nt AUTO -bb 1000 -alrt 1000).

### Annotation of genomes

All genomes included in the reference dataset (651) were annotated using our annotation pipeline to ensure consistency of annotations Annotation_Tables/1_Input/Annotations.txt [38]. Coding sequences were predicted using Prokka v1.14 [46] (settings: --kingdom Archaea -- addgenes --force --increment 10 --compliant --centre UU --cpus 20 --norrna –notrna). For functional annotation of proteins several additional databases were used including COGs [47] (downloaded October 2020), arCOGs [48] (2014 version), KO profiles from the KEGG Automated Annotation Server [49] (downloaded April 2021), the Pfam database [50] (release 34.0), the TIGRFAM database [40] (release 15.0), the Carbohydrate-Active enZymes (CAZy) database [51] (v7, downloaded August 2020), the Transporter Classification Database [52] (downloaded April 2021), the Hydrogenase database [53] (HydDB, downloaded July 2020), and NCBI_nr (downloaded Aug 2021). In addition to this, protein domain predictions were carried out using InterProScan [54] (v5.62-94.0, setting: --iprlookup --goterms).

Annotations for the respective databases were carried out as follows. COGs, arCOGs, KOs, PFAMs, TIGRFAMs, and CAZymes were all identified using hmmsearch v3.1b2 [39] (settings: -E 1e-5). The Transporter Classification Database and Hydrogenase Database were queried using BLASTp v2.7.1 [55] (settings: -evalue 1e-20). For database searches the best hit was selected based on the highest e-value and bit-score and summarised for Caliditerrarchaeota in Supplementary Table 3 and for the full set Annotation_Tables/1_Input/Annotations.txt [38]. Multiple hits were allowed for InterProScan domain annotations using a custom script for parsing results Annotation_Tables/parse_IPRdomains_vs2_GO_2.py [38]. Best blast hits against the NCBI_nr database were identified using DIAMOND v0.9.22..123 [56] (settings: blastp --more-sensitive --evalue 1e-5 --no-self-hits). CRISPR systems were identified in Caliditerrarchaeota genomes using the CRISPRCasTyper v1.8.0 tool [57]. Caliditerrarchaeota genomes were taxonomically classified using GTDB-Tk (setting: –skip_ani_screen)[58].

### Analysis of NADH- [NiFe]- and [FeFe]-hydrogenases

To confirm annotation of putative hydrogenase subunits predicted catalytic subunits from Caliditerrarchaeota genomes were identified and aligned to reference sets of previously published NiFe and FeFe hydrogenase catalytic subunits [59,60] using MAFFT-LINSI v7.407 (settings: reorder). Alignments were trimmed using BMGE-1.12 (settings:-tAA -m BLOSUM30 -h 0.55) and manually screened for presence of conserved active sites in putative Caliditerrarchaeota hydrogenases. Caliditerrarchaeota sequences missing conserved active site residues were removed from the dataset and remaining sequences were realigned with MAFFT-LINSI v7.407 (settings: reorder) and trimmed with BMGE-1.12 (settings:-tAA -m BLOSUM30 -h 0.55). Phylogenetic trees were inferred using IQTree v2.1.2 (settings: -m LG+C20+F+R -nt AUTO -ntmax 25 -bb 1000 -alrt 1000).

### Comparative genomics

Output from the annotation pipeline was used for comparative genomics. Most genomes were clustered into class-level lineages, whilst DPANN and most uncultivated taxa were defined at the level of phylum. Presence/absence patterns of proteins encoded by these clades were determined using a custom R script Annotation_Tables/Count_Tables.r [38]. Briefly, the number of occurrences of each gene in every genome in the dataset was tallied and used to generate summary tables using the ddply function within the plyr (v1.8.8) package. Heatmaps were then plotted by converting the summary tables into presence/absence matrices followed by application of the ddply function to summarize counts across the different phylogenetic clusters. Data was then visualized as a heatmap using ggplot2 (v3.4.2). Genome reconstruction for Supplementary Figure 21 was carried out manually using KEGG annotations. Genes were classified as present (found in more than 50% of genomes, indicated with filled in circle), partially present (found in 0 - 50% of genomes, indicated with half-filled circle), or absent (missing from all genomes) for Supplementary Figure 21.

### Amino acid frequency analysis

To investigate signatures of halophily across our reference set of genomes, amino acid frequencies were calculated using a custom R script Amino_Acid_Analysis/AA_Freq_analysis.R [38]. Briefly, predicted protein sequences from each genome were concatenated and the frequency of each amino acid (i.e. number of its occurrence) was calculated and divided by the total length of the concatenated sequence. Principal components analysis was carried out using prcomp from the stats package (v4.4.0) on a matrix of calculated amino acid frequencies per individual genome across the entire dataset. Statistical significance of amino acid frequency variation between lineages within the reduced dataset was calculated using one-way ANOVA tests from the stats package (v4.4.0) with a Games-Howell Post-hoc evaluation of significance from the rstatix (v0.7.2) package. P-values were used to generate groups with letter designations to simplify representation with multcompview (v0.1-9). Acidity of predicted protein sequences was calculated using the peptides package (v2.4.5) isoelectric point command applied on a protein-by-protein basis to each genome. Calculated pI values were then plotted in density plots on a genome-by-genome basis.

### Gene tree-species tree reconciliation

Gene tree-species tree reconciliation analyses were performed using ALE v1.0 [36,37], ALE/ALE_arCOGs_final.md [38]. Protein clusters were generated using arCOG annotations produced using hmmsearch as described above (section Genome Annotation). To account for gene fusion of arCOGs in some taxa, proteins with more than 50 aa remaining after removal of the sequence part assigned to an arCOG were screened for secondary arCOG annotations and split up, with this procedure being repeated up to four times. Clusters were then cleaned by removing sequences containing Xs prior to alignment. Clusters with 1000 sequences or less were aligned using MAFFT L-INS-I v7.407 (settings: reorder) whilst those with more than 1000 sequences were aligned with MAFFT E-INS-I v7.407 (settings: reorder). All alignments were trimmed using BMGE-1.12 (settings:-tAA -m BLOSUM30 -h 0.55). Clusters with three or fewer sequences were removed from further analysis.

To evaluate gene clusters, initial trees were inferred from trimmed alignments using FastTree v2.1.11 (settings: -lg -gamma) and KEGG annotations mapped to sequences in the tree. A custom script ALE/Split_Count_v2.py [38] was used to screen these phylogenies for long branches and the monophyly of KEGG annotations within trees. Long branches were defined as branches that have lengths greater than 5 times the interquartile range for the tree in which they were found, and corresponding sequences were flagged for removal. Instances where multiple KEGG annotations were assigned to one arCOG cluster were flagged for splitting if more than 10% of sequences in the tree were annotated with the relevant KEGG number and all of these were monophyletic. A total of 424/9481 (4.5%) of clusters were identified as potential candidates for splitting which were then manually reviewed resulting in 74 clusters (0.8%) being split for a total of 9555 individual clusters. Following long branch removal and splitting of clusters, retained sequences were realigned with either MAFFT L-INS-I v7.407(less than or equal to 1000 sequences) or MAFFT E-INS-I v7.407(>1000 sequences) and trimmed with BMGE-1.12 as described above.

Once final alignments were produced guide trees were inferred (settings: -m LG+G+F --score-diff all -nt 1 -wbtl -B 1000 -bnni) and model tests carried out (settings: -m MF -mset LG -madd LG+C10,LG+C10+G,LG+C10+R,LG+C10+F,LG+C10+R+F,LG+C10+G+F,LG+C20,LG+C20 +G,LG+C20+F,LG+C20+G+F,LG+C20+R,LG+C20+R+F --score-diff all -T 1) using IQTree v2.1.2. Model test results were summarised and alignments were distributed into two categories: category one comprised alignments where the best fitting model was a C-series model. For computational feasibility, trees for category 1 alignments were inferred using the rapid PMSF method, which approximates full profile mixture models [61]. Non-mixture models were chosen as best model in category 2 alignments, for which trees could be inferred without PMSF. Trees were inferred for all alignments using IQTree v2.1.2 (settings: -T 1 -wbtl -B 1000 -pers 0.2 -nstop 500) and for category 1 alignments the additional command to run using a PMSF model (-ft Phylogenies/single/iqtree_lg_guide/{1}.treefile).

Following inference of single gene trees, ALE reconciliation was performed for all 9555 arCOG clusters against the species tree inferred using the 50% best marker genes described above. The species tree was rooted between the DPANN and all other archaea for reconciliations, based on previous analyses [4,62]. A first round of reconciliations were performed using ALEm1_undated v1.0^15^ for each gene family including CheckM2 v0.1.3 estimates of genome completeness to correct for incomplete genomes using the fraction_missing flag. In order to carry out ancestral genome reconstructions, the probabilities that gene families originate at the root of the species tree was optimised for each arCOG category by maximum likelihood using a python script ALE/setup_OR_estimation.py [38] and summary statistics extracted using another python script ALE/parse_or.py [38] for each arCOG. This origination model implies that the prior probability of gene family origination is the optimised value, OR, at the root, and (1-OR) divided by the number of non-root nodes elsewhere. Following optimisation of root origination rate, a second run of ALE was performed with ALEm1_undated v1.0 including the maximum likelihood origination rate as a correcting factor with the flag O_R and completeness scores with the flag fraction_missing as above. Inferred gene statistics were parsed using a custom python script (ale_parser.py). Data was filtered for nodes of interest using another custom python script (Ale_summary.py) and genome reconstruction carried out manually. arCOGs were classified as likely present (PP >= 0.8, filled in circle), possible present (0.8 > PP >= 0.5, half-filled circle), or likely absent (PP < 0.5, empty circle) for genome reconstructions.

### Sisterhood Analysis

To investigate occurrences of sisterhood relationships between archaeal taxa, single gene trees produced for ALE analyses were processed using a custom python script HGT_Analysis/Tree_Analyser.py [38]. Briefly, GTDB taxonomy was mapped to leaves in the trees and for all Order and Family level lineages the relative frequency of sisterhood was calculated per tree. To calculate relative frequency of sisterhood each instance of the target lineage in the tree was identified and progressive searches conducted of nodes upstream until a member of a different lineage was found. Maximum branch length (3) and minimum branch support (0.7) cutoffs were applied and sisterhood frequencies were not calculated for trees in which length or support values fell outside these cutoffs. Once a sister clade was identified the relative frequency of different lineages within the sister clade was calculated per tree. Following this the relative frequencies across all trees for each lineage were summed together. Plots were then produced using a custom R script HGT_Analysis/Sisterhood_Analyses.R [38].

## Results and discussion

### Caliditerrarchaeota form a thermophilic, non-halophilic sister phylum of the Nanohaloarchaeota

To assess the phylogenetic relationships among DPANN cluster 2 lineages, we built upon a previously established approach [4] to identify a set of universal single copy marker genes least affected by HGT (25% and 50% top ranked markers). All concatenated phylogenies resolved a monophyletic clade within the DPANN consisting of Nanohaloarchaeota, Aenigmatarchaeota, and Caliditerrarchaeota (see Classification of Caliditerrarchaeota) (Figure 1, Supplementary Figures 1 - 4). Caliditerrarchaeota were recovered as the sister lineage of the Nanohaloarchaeota in phylogenies inferred using the 25 and 50% top marker genes (100/100 and 99.8/100 SH-like aLRT/ultrafast bootstrap support, respectively) (Figure 1, Supplementary Figures 1 and 2). Their placement sister to both Nanohaloarchaeota and the Aenigmatarchaeota lineage PWEA01 (recently proposed as Haloaenigmatarchaeaceae [18]) (26/69 and 89.2/95 SH-like aLRT/ultrafast bootstrap support respectively) in phylogenies using the 25 and 50% worst makers (Supplementary Figures 3 and 4) is likely an artifact of incongruent markers. Together, this suggests that the Nanohaloarchaeota and Calditerrarchaeota branch with the Aenigmatarchaeota, consistent with previous analyses [18], and form a clade which we here refer to as ACN (for Aenigmatarchaeota, Calditerrarchaeota and Nanohaloarchaeota).

**Figure 1.**
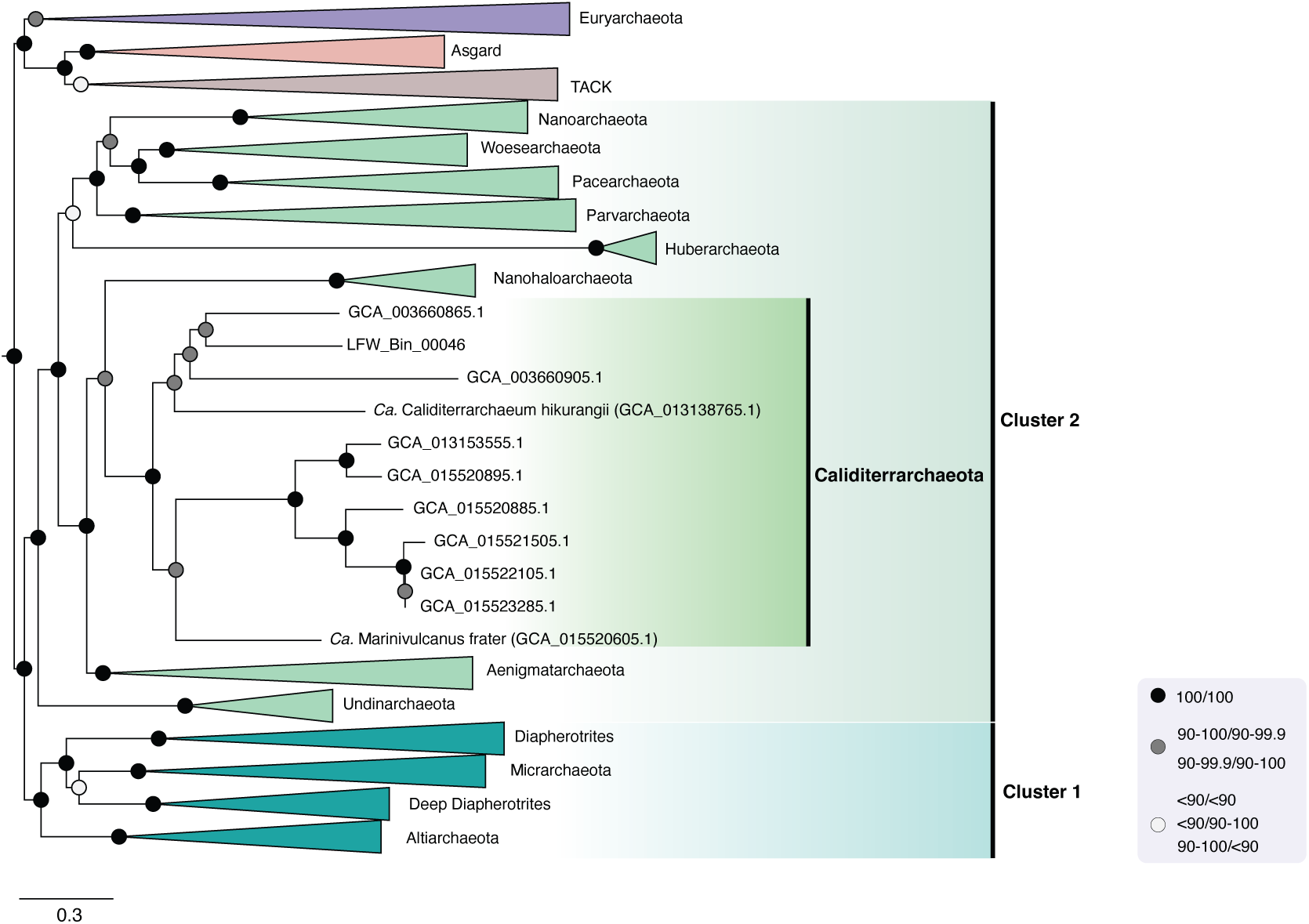
Phylogenetic tree showing placement of Caliditerrarchaeota based on a concatenated alignment of 50% top ranked marker genes and the 651 species set. The alignment was trimmed with BMGE (Alignment length = 12,397 aa). A Maximum Likelihood (ML) phylogenetic tree was inferred using IQ-Tree with the LG+C60+F+R model and ultrafast bootstrap approximation (left) and SH-like approximate likelihood tests (right), each run with 1000 replicates. The tree has been artificially rooted between DPANN and other Archaea. The full uncollapsed version of the tree is provided as Supplementary Figure 1. Scale bar: Average number of substitutions per site.

Removal of the 10, 30, or 50% fastest evolving [63] (Supplementary Figures 5 - 10) or compositionally biased [64] (Supplementary Figures 11 - 16) sites had little to no impact on the placement of the Caliditerrarchaeota in either of the top 25 or 50% marker gene phylogenies. Likewise, addition of genomes from the recently described *Asbonarchaeaceae* lineage [22] (a sister lineage to the *Nanosalinaceae* within the Nanohaloarchaeota) further supported the sister relationship of Caliditerrarchaeota and Nanohaloarchaeota (Supplementary Figs 17 - 20).

### Caliditerrarchaeota are putative non-halophilic symbionts

To provide insights into the metabolism and salt adaptation of Caliditerrarchaeota, we analysed 11 publicly available MAGs (genome completeness and contamination ranging from 64 – 91% and 0 – 2.19%, respectively) (Supplementary Tables 1 and 2) derived from hydrothermal vent [32] and deposit [63], methane seep, and radioactive site metagenomes [64]. Only one previous study investigated Caliditerrarchaeota metabolism, concluding that they are fermentative symbionts with a higher than typical number of transporters compared to other DPANN based on a single MAG (GCA_902384675.1) [64].

Metabolic predictions revealed that the 11 representatives of the Calditerrarchaeota have limited catabolic and anabolic potential similar to other DPANN archaea (Supplementary Table 5, Supplementary Figures 21 - 25, Supplementary Information). Specifically, while encoding a glycolytic pathway and enzymes for the degradation of complex carbohydrates, most representatives lack TCA cycles genes and genes for any of the known carbon fixation pathways of archaea [65]. Many biosynthetic pathways such as for amino acids, vitamins, purine and pyrimidine, and archaeal lipids are incomplete. However, similar to other DPANN, representatives encode the capacity for interconversion of nucleotides as well as ribose 1,5- bisphosphate isomerase (K18237, 8 MAGs) and ribulose bisphosphate carboxylase (RbcL; RuBisCo, K01601, 7 MAGs), which are hypothesized to function in the salvage of nucleosides [4]. However, all Caliditerrarchaeota do encode a complete A/V-type ATP synthase, and some representatives encode Group A [FeFe]-hydrogenases (2 MAGs), a group 3 [NiFe]- hydrogenase (2 MAGs), putative Group 4 [NiFe]-hydrogenase (3 MAGs) and a putative NADH-dehydrogenase (Supplementary Text, Supplementary Tables 3 and 6, Supplementary Figures 26 - 28). The presence of reverse gyrase homologs in all but one Caliditerrachaeota MAG (GCA_902384675.1) is consistent with their distribution in high-temperature marine sediments.

PCA analyses of amino acid frequencies, isoelectric point (pI) profiles and amino acid frequencies inferred for all representatives in our taxon set, revealed that Caliditerrarchaeota are not extreme halophiles. Specifically their proteomes clustered with non-halophilic Aenigmatarchaeota (Figure 2a) and isoelectric point (pI) profiles showed no signs of increased proteome acidity characteristic for halophilic Archaea (Figure 2b). Further, we did not observe statistically significant variation in individual amino acid frequencies for Caliditerrarchaeota except for tryptophan and proline (Figure 2c, Supplementary Table 7). However, these analyses revealed a subclade within the Aenigmatarchaeota with hallmarks of adaptations to higher salt conditions (Figure 2a). Isoelectric profiles of PWEA01 protein coding sequences also showed signs of a shift towards higher acidity in the proteome, and statistically significant variation in amino acid frequencies were observed for nine amino acids, including aspartate and glutamate, which are typically elevated in halophiles [19,66]. Other amino acids that significantly vary in frequency amongst Nanohaloarchaeota, such as cysteine and serine, do not differ in PWEA01. Overall, the amino acid profiles of the PWEA01 clade appear to occupy an intermediate position between Nanohaloarchaeota and other archaea in PCA plots. This indicates that members of this group are halophiles, which is in agreement with the recent discovery of PWEA01 representatives, now also referred to as Haloaenigmatarchaeaceae, in hypersaline brines from the Danakil Depression [18].

**Figure 2.**
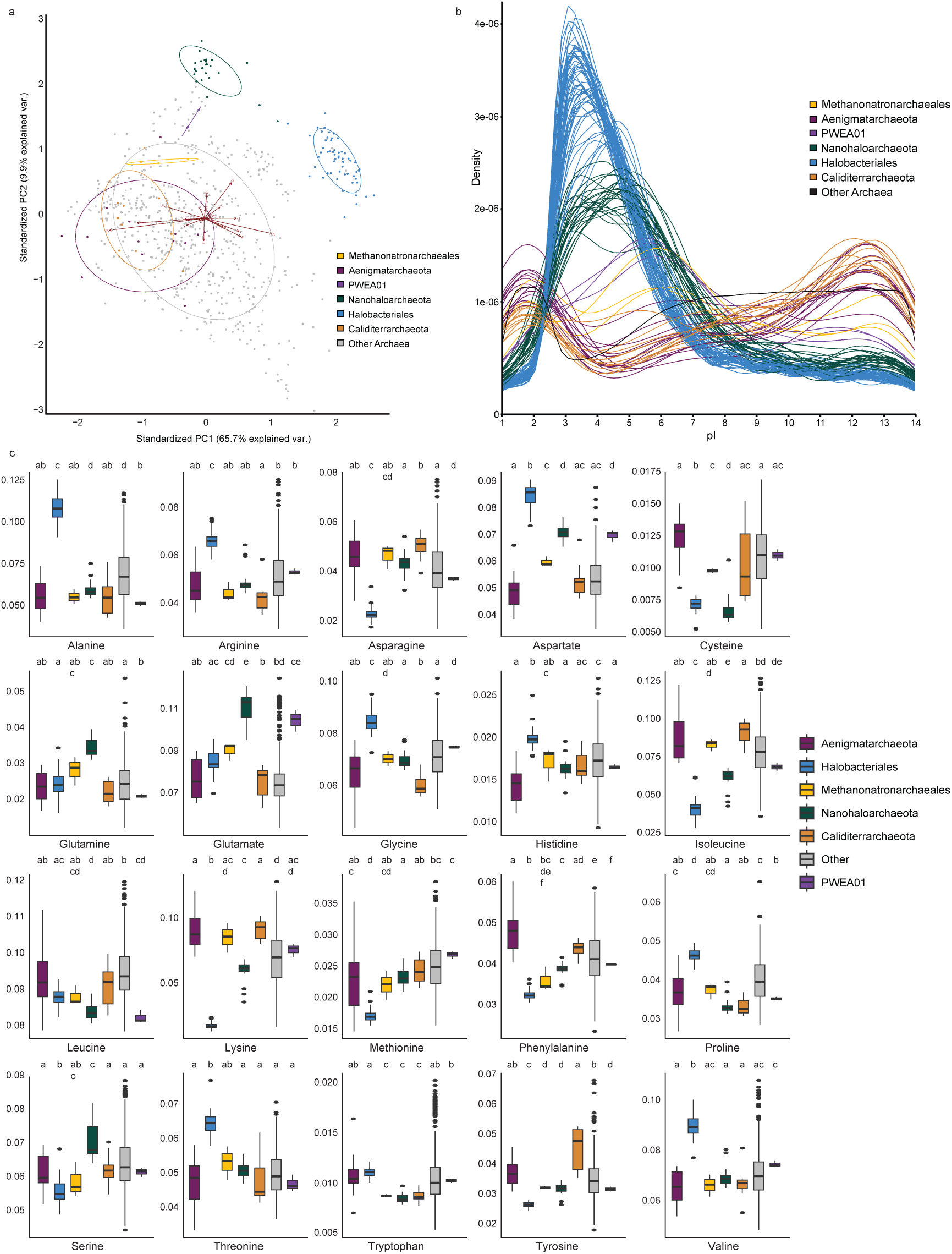
Amino acid composition of both ACN and known halophilic lineages. **a)** PCA showing distribution of genomes in the 651 genome dataset based on amino acid profile. **b)** Protein isoelectric point frequency plot of genomes in the 651 genome dataset. Plots show isoelectric points per genome with genomes colour-coded according to which lineage they are assigned to. Genomes assigned to lineages outside the ACN (Aenigmatarchaeota, Caliditerrarchaeota, and Nanohaloarchaeota) cluster or known halophiles are grouped into a single line for readability **c)** Boxplot showing individual amino acid frequencies with statistical significance tests across the 651 genome dataset. For all plots ACN lineages and known halophiles are highlighted with colour whilst all other lineages are shown in grey. Letters above plots indicate statistically significant differences determined using one-way ANOVA test and a Games Howell Post-hoc evaluation of significance and then simplified with letter codes using multcompview.

### The evolution of metabolic gene repertoires and halophilic lifestyles in the ACN clade

To elucidate the evolutionary history of gene repertoires in the ACN clade and the adaptation to halophilic lifestyles in some member lineages, we next performed gene tree-species tree reconciliations using Amalgamated likelihood analysis (ALE) [36,37,67]. We rooted our species tree between DPANN and the rest of the archaea in agreement with several recent publications [4,62,68]. Alternative root positions, such as within the former ‘Euryarchaeota’ [69] or between ‘Euryarchaeota and TACK/Asgard Archaea’ [27,70], are unlikely to significantly impact our results because the ACN lineage represents a derived clade relative to any of the proposed root positions. Ancestral genome content was inferred based on posterior probabilities (PP) of arCOG family presence in any node of interest (Supplementary Table 8). We considered gene families with PP values between 0.5-0.8 to have low support and greater than 0.8 to be of moderate support for having been present at the nodes examined.

Reconstructed metabolic processes support an ancestral fermentative lifestyle for all last common ancestors examined; i.e. Aenigmatarchaeota (LCA-A), Caliditerrarchaeota (LCA-C), Nanohaloarchaeota (LCA-N), Caliditerrarchaeota-Nanohaloarchaeota (LCA-CN), Aenigmatarchaeota-Caliditerrarcheota-Nanohaloarchaeota (LCA-ACN) with acetate the likely end-product for all ancestors except for the Nanohaloarchaeota and Haloaenigmatarchaeaceae. Specifically, all ancestors were predicted to encode the majority of proteins for glycolysis but to lack respiratory complexes and various genes for TCA cycle enzymes (Figure 3, Supplementary Table 8). Citrate synthase, either encoded by two (K22224 and K01905) or a single gene (K01647), is the only TCA cycle protein inferred to be potentially present in most ACN ancestors (LCA-A: PP=0.97/1/0.04; LCA-C: PP=0.85/1/0.07; LCA-N: PP=0.39/0.28/0.93; LCA-CN: PP=0.84/0.9/0.56; LCA-ACN: PP=0.95/1/0.45) and citrate may therefore represent an important intermediary in the central carbon metabolism in these archaea. Nanohaloarchaeota appear to have had an ancestral citrate synthase (K22224, K01905), replaced by an alternative version (K01647), with both types potentially present in the common ancestor of Nanohaloarchaeota and Caliditerrarchaeota. Given the absence of other TCA cycle genes and the potential reversibility of the reaction carried out by these enzymes it is possible that these DPANN are instead converting citrate into pyruvate enabling either ATP production via fermentation to acetate or carbon storage via gluconeogenesis as previously suggested for Nanohaloarchaeota [11]. The predicted presence of a putative tri-carboxylic acid importer capable of citrate import in all ancestors examined (arCOG04469: LCA-A: PP=1; LCA-C: PP=1; LCA-N: PP=0.98, LCA-CN: PP=1, LCA-ACN: PP=1, LCA-P: PP=1; LCA-PB: PP=1) could provide the means for uptake of exogenous citrate.

**Figure 3.**
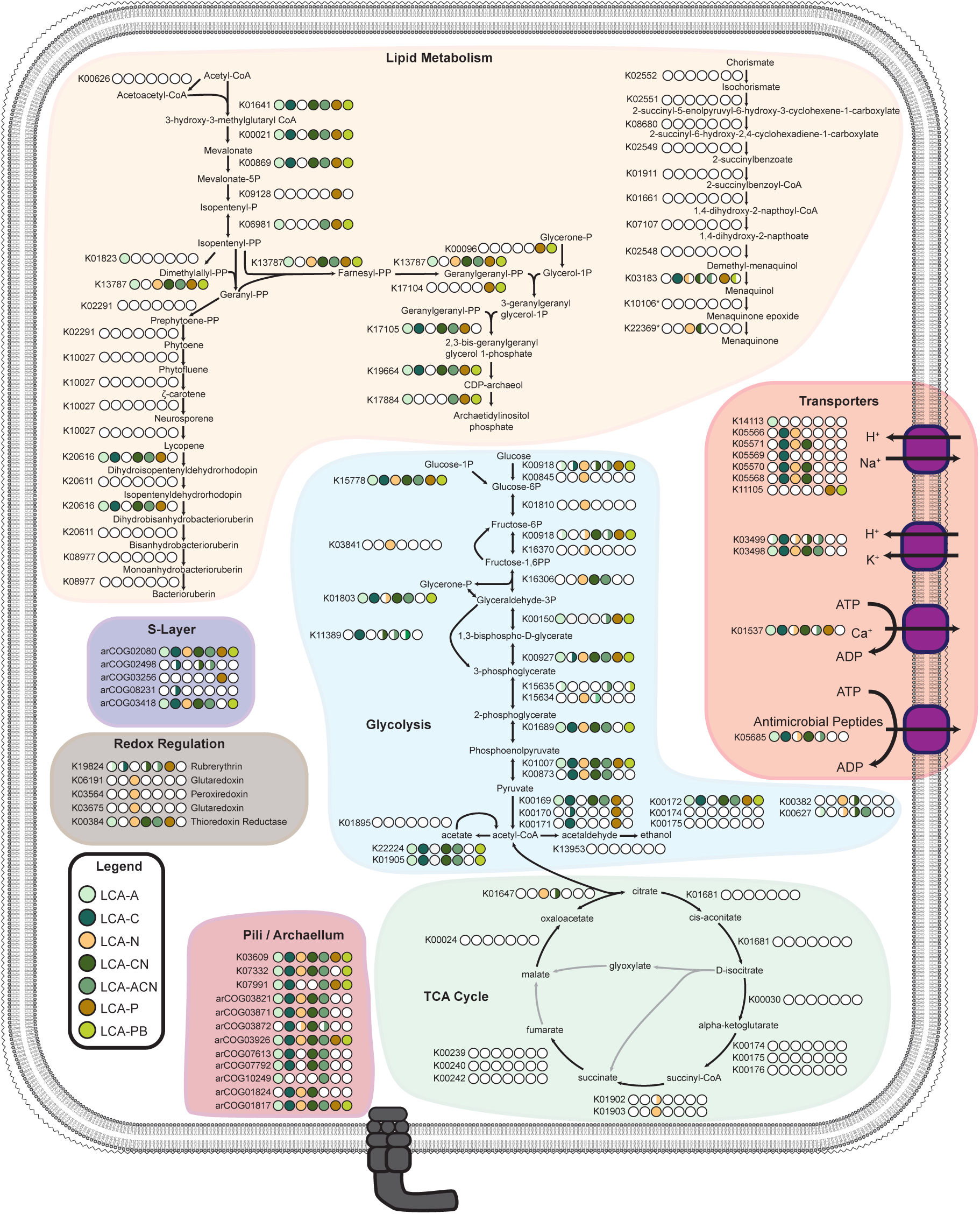
Ancestral genome reconstruction for ACN cluster ancestors. Figure shows predicted presence/absence of major metabolic/symbiosis associated genes in various ACN lineage ancestors as predicted from ALE gene tree/species tree reconciliation. Half circles indicate a posterior probability for gene presence of ≥0.5 <0.8 and full circles indicate a posterior probability of ≥0.8. The data used to produce this figure is provided in Supplementary Table 7.

All ancestors examined, with the exception of LCA-N, were inferred to have complete or near-complete phospholipid biosynthesis pathways allowing production of archaeol from acetyl-coA indicating their capability of synthesising membrane lipids (Figure 3, Supplementary Table 8). In contrast, the majority of quinone and carotenoid biosynthesis pathways were predicted to be absent from all ancestral genomes (Figure 3). Absence of quinone synthesis pathways is unsurprising considering that most Cluster 2 DPANN are generally thought to lack respiratory complexes that require quinones.

The analysed ancestors were also predicted to encode pili/archaellum subunits, although individual KOs and arCOGs showed differential distributions across the various lineages (Figure 3, Supplementary Table 8). Typically, ancestral proteomes comprise the scaffold proteins that are also identified in genomes of extant representatives, whilst the accessory proteins, including the pilins themselves, display a variable distribution (Figure 3). Pilins may be involved in symbiont-host interactions [71–73] with specific pilins potentially allowing interaction with specific hosts. However, the low number of cultivated DPANN and known hosts currently limits experimental validation. A minimum of two S-layer proteins were predicted to be encoded by all ancestral genomes with all lineages except Haloaenigmatarchaeaceae predicted to share the same two arCOGs (arCOG02080: LCA-A: PP=0.91; LCA-C: PP=1; LCA-N: PP=1, LCA-CN: PP=1, LCA-ACN: PP=0.98, and arCOG03418: LCA-A: PP=1; LCA-C: PP=1; LCA-N: PP=0.93, LCA-CN: PP=1, LCA-ACN: PP=1). The Haloaenigmatarchaeaceae ancestor was instead predicted to encode an alternative S-layer protein, assigned to arCOG03256 (PP=1) rather than arCOG03418 (PP=0.35). Subunits of the sodium-hydrogen antiporter (arCOG03076, arCOG03079, arCOG03082, arCOG03099, arCOG03121, arCOG01537, arCOG01962), which is particularly important in regulating osmotic balance in extreme halophiles, was predicted to be present in all individual ancestors of the various ACN lineages but not in LCA-ACN, with LCA-A and LCA-CN predicted to encode different KOs (Supplementary Table 8). Absence of these genes from LCA-ACN may indicate that each lineage acquired their antiporter complexes via HGT. Consistent with this scenario, single gene trees resolved Nanohaloarchaeota sequences as either sister to or branching within halobacterial clades for all antiporter subunits whilst Aenigmarchaeota and Caliditerrarchaeota sequences typically form monophyletic clades sister to Thermoplasmatota. However, it is also possible that the amino acid biases associated with adaptation to high salt conditions has caused artificial attraction of the nanohaloarchaeal and halobacterial sequences. LCA-N was predicted to encode at least four different redox regulation proteins including two glutaredoxins (arCOG02608: PP=0.94 and arCOG01297: PP=0.98), one peroxiredoxin (arCOG00314: PP=0.88) and a thioredoxin reductase (arCOG01296: PP=0.97) but not rubrerythrin. None of these protein families have high posterior probabilities for being present in the proteomes of any of the ancestors, though there is moderate support for rubrerythrin in some of them (K19824: LCA-A: PP=0.55; LCA-C: PP=0.54; LCA-N: PP=0.11, LCA-CN: PP=0.50, LCA-ACN: PP=0.55, Figure 3). The absence of rubrerythrin from LCA-N is consistent with a transition into an oxic environment alongside their haloarchaeal hosts whilst the putative presence of it in some of the other lineages may indicate their adaptation to an anoxic environment. LCA-N is the only DPANN ancestor predicted to encode both photolyase (arCOG02840: LCA-A: PP=0.04; LCA-C: PP=0.07; LCA-N: PP=0.99, LCA-CN: PP=0.49, LCA-ACN: PP=0.28) and RecB (arCOG00802: LCA-A: PP=0.12; LCA-C: PP=0.06; LCA-N: PP=0.99, LCA-CN: PP=0.47, LCA-ACN: PP=0.27) which both may play a role in DNA repair. This is particularly relevant as many hypersaline systems cause exposure to high UV-radiation [74] necessitating adaptation to UV-induced DNA damage.

Finally, we analysed the ALE output to infer copy number change along branches of interest (Figure 4, Supplementary Table 9). While the Nanohaloarchaeota and Haloaenigmatarchaeaceae seem to have distinct protein families with multiple copies per proteome, the Kef K^+^ transporter (arCOG01955) has the highest copy number increase along the branches leading to both lineages. The Kef-type potassium transporters play an important role in halophiles that use a salt-in strategy by facilitating the uptake of K^+^ ions allowing to maintain osmotic balance whereas Na^+^ and Mg^2+^ ions are excluded from the cytoplasm [66]. Acquisition of multiple copies of the Kef-type transporter may have allowed members of both the Nanohaloarchaeota and Haloaenigmatarchaeaceae to adjust to hypersaline systems in the absence of additional protective measures (e.g. osmoprotectants such as glycine betaine) that are generally present in *Halobacteriales* [75].

**Figure 4.**
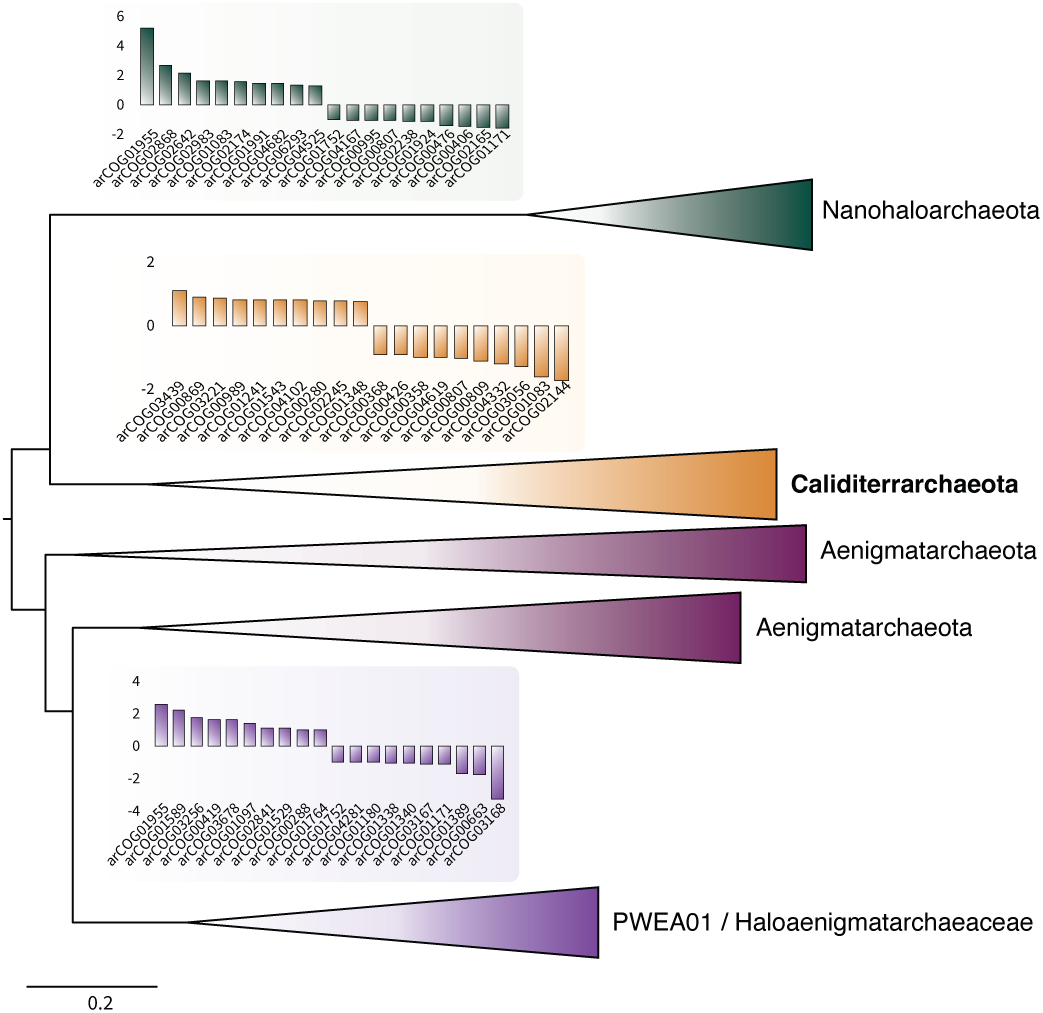
Predicted gene copy number changes along branches of interest within the ACN cluster. Figure shows the 10 genes with the highest copy number increase and the 10 genes with the highest copy number decrease predicted from ALE gene tree/species tree reconciliations along branches of interest within the ACN cluster. Data used to produce this figure is provided in Supplementary Table 8.

### Who are the hosts of uncultivated representatives of the ACN lineages?

It has previously been shown that some DPANN archaea exchange genes more frequently with their hosts than with other taxa, such as Nanohaloarchaeota with Halobacteriales, Micrarchaeota and Parvarchaeota with Thermoplasmatales, Huberarchaeales with Altiarchaeales, and Nanoarchaeales with Sulfolobales [4]. To investigate potential hosts of the Caliditerrarchaeota and Haloaenigmatarchaeaceae lineages, we calculated the relative frequency of sisterhood relationships for all trees used in the ALE analysis at the level of Order and Family from the GTDB taxonomy (Figure 5, Supplementary Table 10 and 11, Supplementary Figure 29). While we could not identify a clear signal for Caliditerrarchaeota, Haloaenigmatarchaeaceae representatives displayed an increased frequency of sisterhoods with PWKY01 (Halarchaeoplasmatales), suggesting that members of the PWKY01 may interact with or serve as hosts for some representatives of the Haloaenigmatarchaeaceae clade.

**Figure 5.**
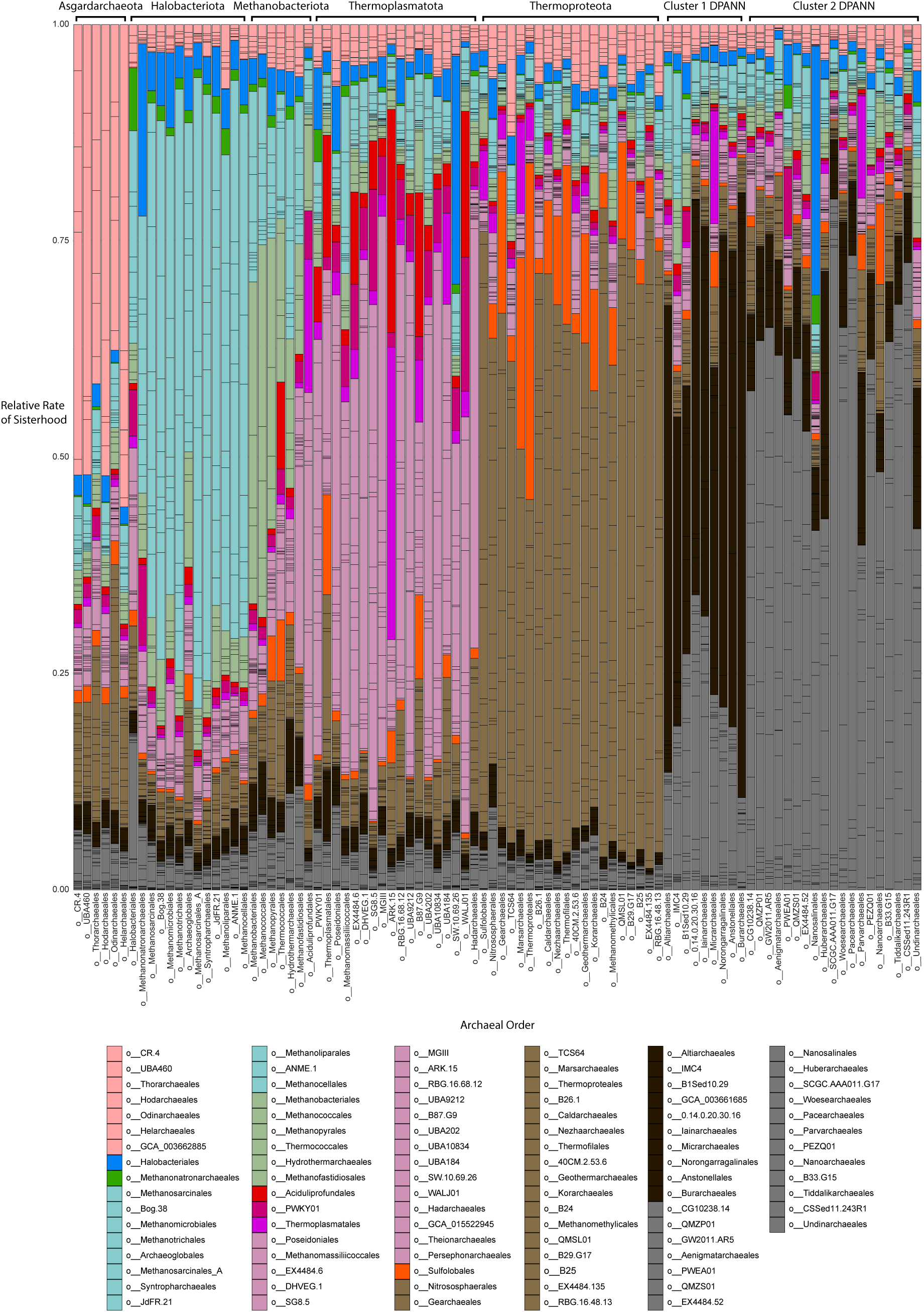
Rate of sisterhood for GTDB order level lineages across 9,555 gene trees used for ALE analysis. Relative sisterhood frequencies were calculated per gene and summed for each taxonomic group. Related lineages have been clustered together using colours and specific lineages that show higher rates of sisterhood with DPANN lineages highlighted with unique colours. The data used to generate this plot is available in Supplementary Table 10.

It is conspicuous that all DPANN symbiont-host partnerships with observable increase in sisterhood rates (except Huberarchaeales and Altiarchaeales) occur in extreme environments (hypersaline, acidic, and/or high-temperature). It seems possible that the selective pressures involved in adapting to these environments favours the acquisition of genes related to the specific extremophily from the host at rates higher than those seen in other DPANN symbiont-host partnerships from less extreme environments. Consistent with this, several copies of the KefB Potassium-efflux system gene (the highest increase in copy number for both Nanohaloarchaeota and Haloaenigmatarchaeaceae) from DPANN halophile genomes placed as sister to their respective/putative hosts (*Halobacteriales* and PWKY01) in single gene trees ALE/Treefiles [38]. Alternatively, given that extreme environments typically support less diverse microbial communities, symbiont-host gene transfers may be more easily detected from background HGT. Additionally, amino acid composition biases, such as those displayed by extreme halophiles and thermophiles, are known to cause compositional attraction artefacts in trees, particularly when limited signal is available as is the case for single gene trees. Thus, a fraction of sisterhood relationships between extremophilic DPANN and their putative hosts may be the result of erroneous gene trees rather than genuine gene transfer [4,22]. Finally, we note that the trees used for this analysis were limited to archaeal taxa only. However, many DPANN lineages occupy environments dominated by Bacteria and some representatives have been proposed to associate with bacterial hosts [4,63,76,77]. The absence of bacterial sequences from our trees due to computational limitations of the gene tree-species tree reconciliation workflow, limit the confidence with which sisterhood relationships can be determined. Therefore whilst our results suggest a possible relationship between the Haloaenigmatarchaeaceae and the Halarchaeoplasmatales which remains to be confirmed experimentally, they do not allow us to identify hosts for DPANN that do not interact with other Archaea.

### Classification of Caliditerrarchaeota

Classification of the Caliditerrarchaeota genomes with GTDB-Tk [58] indicated that all 11 MAGs included in this analysis belong to the existing candidate phylum referred to as EX4484-52. The included genomes represent 6 families, 7 genera, and 10 species. Consistent with the SeqCode Registry requirements for designation of type material we designate *Candidatus* Caliditerrarchaeum hikurangii (Assembly ID: GCA_013138765.1, Estimated completeness ∼90%, Estimated contamination: 1.6%, tRNAs: 20/20, 5S and partial 16S rRNA genes present) as type material for the phylum EX4484-52 with the new proposed name Caliditerrarchaeota, and the corresponding family (Caliditerrarchaeaceae), order (Caliditerrarchaeales), and class (Caliditerrarchaeia), following the principle of propagation from type material as outlined in SeqCode (Section 3, Rule 15). Additionally, we designate *Candidatus* Marinivulcanus frater (Assembly ID: GCA_015520605.1, Estimated completeness: ∼91%, Estimated contamination: 0%, tRNAs: 15/20, 5S rRNA gene present) as type material for the novel family Marinivulcanaceae.

Description of ‘*Candidatus* Caliditerrarchaeum’ gen. nov.

‘Caliditerrarchaeum’ (Cal.i.di.terr.ar.chae’um. L. adj. *calidus* warm, hot: L. fem. n. *terra* earth: N.L. neut. n. *archaeum* archaeon from Gr. adj. *archaios −ê −on* ancient; N.L. neut. n. An archaeon that inhabits hot earth, reflecting the preference for this lineage to inhabit hot sediments)

Description of ‘*Candidatus* Caliditerrarchaeum hikurangii’ sp. nov.

‘Caliditerrarchaeum hikurangii’ (hi.ku.ran.gi’i N.L. neut. n. *hikurangii*, of Hikurangi, due to the sample site from which the genome originates being found in the Hikurangi Margin, an active subduction zone in the deep sea located off the east coast of New Zealand)

Description of ‘*Candidatus* Marinivulcanus’ gen. nov.

*‘Candidatus* Marinivulcanus’ (mar.in.ni.vul.kan.nus. L. adj. *marīnus* marine: L. masc. n. *vulcānus* fire; N.L. masc. n. marine volcano, referring to the type genome within this genus originating from marine volcano sediments)

Description of ‘*Candidatus* Marinivulcanus frater’ sp. nov.

Marinivulcanus frater (fra.ter. L. masc. n. *frater* brother; referring to the site this genome originates from, Brothers Volcano, a submarine volcano located in the Pacific Ocean)

### Concluding Remarks

The Caliditerrarchaeota, which we infer to be fermentative thermophiles, occupy an evolutionarily important position as the closest sister to the extremely halophilic Nanohaloarchaeota (including both *Nanosalinaceae* and *Asbonarchaeaceae*), but are not themselves halophiles. The inclusion of Caliditerrarchaeota in gene content analyses revealed that heightened tolerance to DNA damage and increased capacity for import and export of salt ions were key in the evolution of all currently known halophilic DPANN archaea. However, the results also indicated important differences between lineages such as the adaptation to oxic environments in the Nanohaloarchaeota and preferences for different protein families involved in the same functions e.g. S-layer proteins, Na^+^/H^+^ antiporter, and citrate metabolism enzymes. Furthermore, the predicted acquisition of genes from host species via HGT highlights the potential importance of the symbiotic interactions these DPANN engage in for their adaptation to hypersalinity, although it remains possible these analyses are impacted by compositional attraction artefacts. Future work is necessary to confirm the potential partnership between Haloaenigmatarchaeaceae and Halarchaeoplasmatales and to assess the degree to which compositional biases impact the predicted rates of HGT between halophilic archaea. However, our work provides a framework for investigating the adaptation to hypersalinity in the DPANN archaea that can serve as a foundation for future works examining the evolution halophily across the archaeal domain.

## Acknowledgements

This project has received funding by the Swedish Research Council (VR starting grant 2016-03559 to AS), the Netherlands Organization for Scientific Research Dutch Research Council (NWO) (WISE fellowship to AS and OCENW.M.22.117 to A.S.)), the Moore–Simons Project on the Origin of the Eukaryotic Cell, (Simons Foundation 735929LPI to A.S. as Co-Pi; Gordon and Betty Moore Foundation, GBMF9741 to T.A.W. and A.S.), and the European Research Council (ERC) under the European Union’s Horizon 2020 research and innovation programme (grant agreement No. 947317, ASymbEL to A.S.). Our research is funded by the John Templeton Foundation (63451 to T.A.W. and A.S.; the opinions expressed in this publication are those of the authors and do not necessarily reflect the views of the John Templeton Foundation.

## Author Contributions

JNH and AS conceived the study. JNH, ND, LEVA, CG, TAW, and AS performed genomic and phylogenetic analyses. JNH, TAW, LEVA, CG, and AS conducted data interpretation. JNH, ND, LEVA, CG, TAW, and AS wrote the manuscript. All authors have read and approved the manuscript submission.

## Competing Interests

The authors declare no competing interests.

## Code availability

All custom scripts and workflows used to generate data can be found in our data repository at Zenodo [10.5281/zenodo.14627918] [38]

## Data availability

All datasets generated and/or analysed during this study are available in our data repository at Zenodo [10.5281/zenodo.14627918] [38]. Public databases used in this study are the following: the arCOG database (version from 2014) downloaded from [ftp://ftp.ncbi.nih.gov/pub/wolf/COGs/arCOG/], the KO profiles downloaded from the KEGG Automatic Annotation Server in April 2019 [https://www.genome.jp/tools/kofamkoala/], the Pfam database (Release 31.0) [ftp://ftp.ebi.ac.uk/pub/databases/Pfam/releases/], the TIGRFAM database (Release 15.0) [ftp://ftp.jcvi.org/pub/data/TIGRFAMs/], the Carbohydrate-Active enZymes (CAZy) database downloaded from dbCAN2 in September 2019 [http://bcb.unl.edu/dbCAN2/download/], the MEROPs database (Release 12.0) [https://www.ebi.ac.uk/merops/download_list.shtml], the Transporter Classification Database(TCDB) downloaded in November 2018 [http://www.tcdb.org/download.php], the hydrogenase database (HydDB) downloaded in November 2018 [https://services.birc.au.dk/hyddb/browser/], and NCBI_nr downloaded in November 2018 [ftp://ftp.ncbi.nlm.nih.gov/blast/db/].

## Notes

https://zenodo.org/records/14627918

### Competing Interest Statement

The authors have declared no competing interest.

