## Supplementary material for "Caliditerrarchaeota, a new sister to Nanohaloarchaeota, provides insights into the evolution of DPANN halophily"

**Contents**

**Supplementary Text**

| | |
|---|---|
| **Metabolism of Caldiditerrarchaeota** | **3 - 4** |
| **References** | **5** |
| **Supplementary Figures** | **6 - 34** |

### Supplementary Material

#### Metabolic potential of Calditerrarchaeota

While most Caldíterrarcaheota encode a complete Embden-Meyerhof glycolytic pathway, few TCA cycle genes seem to be present and acetate is likely the end-product of the central carbon metabolism (Supplementary Tables 3 - 5, Supplementary Figure 21), in agreement with inferences from previous work [1]. Like Nanohaloarchaeota, the Calditerrarchaeota genomes also have the capacity for degradation of polysaccharides (potentially Alpha-D-Glucans) with the possibility for products to be fed into the glycolytic pathway. Additionally, Calditerrarchaeota have the capacity for conversion of ribose to fructose via ribulose and arabino-hexulose and subsequent degradation via glycolysis (Supplementary Table 4, Supplementary Figure 21). However, in spite of their limited metabolic capabilities, all Calditerrarchaeota representatives encode a complete A/V-type ATP synthase (Supplementary Tables 3 - 5).

Several Calditerrarchaeota genomes may have the potential to metabolise molecular hydrogen ($H_2$). Specifically, several (36%) Calditerrarchaeota genomes encode Group A [FeFe]-hydrogenases and phylogenetic analysis of two of these subunits supported a placement within the Group A clade (Supplementary Figure 26). In DPANN archaea, these Group A [FeFe]-hydrogenases were recently proposed to re-oxidise ferredoxin potentially reduced during fermentation, resulting in production of $H_2$ [2]. Additionally, five [NiFe]-hydrogenases were identified with the respective large subunit domain proteins assigned to Group 3 (2 cases), NADH dehydrogenases (2) and Group 4 (1 case) hydrogenase, respectively, based on phylogenetic analyses as well as HydDB classification (Supplementary Table 3, Supplementary Table 6, Supplementary Figures 27 - 28). However, the gene neighbourhood of the large subunits of the Group 3 [NiFe]-hydrogenases did not provide information regarding functional context. In contrast, the large subunits of the NADH dehydrogenases that clade with other DPANN archaeal nuoD homologs (Supplementary Figure 27) are part of gene clusters comprising other NADH dehydrogenase subunits except for the electron transfer module NuoEFG, similar to what has previously been observed in some archaeal lineages [3]. The function of these NADH-dehydrogenases and the putative Group 4 [NiFe]-hydrogenase in GCA_902384675.1 remain to be elucidated.

Interestingly, at least some Calditerrarchaeota genomes appear to encode a partial archaeal mevalonate pathway for the formation of archaeal phospholipids (Supplementary Tables 3 - 5, Supplementary Figures 21 and 23). Genes encoding the synthesis of archaeol and archaetidyl-inositol phosphate from geranylgeranyl-PP and glycerol-1 phosphate (K17104, K17105, K19664, and K17884, Supplementary Figures 21 and 23, Supplementary Tables 3 - 5) are present in ~10% - 40% of Calditerrarchaeota genomes analysed. Pathways for biosynthesis of carotenoids and quinones are absent in all Calditerrarchaeota genomes (Supplementary Tables 3 - 5). Furthermore, the Calditerrarchaeota genomes encode a range of amino acid related metabolic pathways with the capacity for biosynthesis/conversion of nine amino acids including a partial shikimate pathway (Supplementary Tables 3 - 5, Supplementary Figure 21). In addition to this, we could identify partial purine and pyrimidine biosynthesis pathways and the capacity for interconversion of nucleotides as well as ribose 1,5-bisphosphate isomerase (K18237) and ribulose bisphosphate carboxylase (RbcL; RuBisCo, K01601) consistent with previous reports of RuBisCo homologs in diverse DPANN lineages likely used for salvage of nucleosides [4]. Yet, various biosynthesis pathways for amino acids, vitamins, and nucleotides are incomplete, suggesting that members of the Calditerrarchaeota are dependent on supplementation via exogenous sources of these compounds.

Calditerrarchaeota genomes encode both DNA Polymerase B3 and D genes but, unlike Nanohaloarchaeota, do not appear to possess a Polymerase B2 gene (Supplementary Tables 3 and 4, Supplementary Figure 24). Their ribosomes share features with those of Nanohaloarchaeota such as the presence of ribosomal protein L40E (absent in Aenigmatarchaeota) and others with Aenigmatarchaeota including the presence of S26 and S28E/S33 (absent in Nanohaloarchaeota) (Supplementary Figure 25). In contrast to both Nanohaloarchaeota and Aenigmatarchaeota, Calditerrarchaeota encode both reverse gyrase (55% of genomes) and DNA gyrase subunit A (55% of genomes) and subunit B (64% of genomes) (Supplementary Figure 24). Considering that all Calditerrarchaeota MAGs except one (GCA_902384675.1) originate from high-temperature marine sediments, it is possible that the absence of reverse gyrase from some representatives is due to MAG incompleteness (Supplementary Table 1). Conversely, GCA_902384675.1, which originates from a moderate temperature radioactive site [1], does not encode any reverse gyrase or DNA gyrase subunits.

Calditerrarchaeota MAGs encode multiple archaellum associated genes (K07332, K07333, K02656, K07329, K07991, K07330, K02650, K02655, K03411 and K07325) with an average of 3.4 copies of K07332 and 5.45 copies of K07333 (Supplementary Figure 21, Supplementary Tables 3 - 5). However, like Nanohaloarchaeota [5] these subunits (predominantly FlaJ, FlaI, FlaF, and FlaK) seem insufficient to constitute complete archaellum apparati due to absence of key subunits such as FlaC, FlaD, and FlaH from all Calditerrarchaeota genomes and absence of FlaG from most (present in 18% of Calditerrarchaeota genomes analysed). Of these subunits, FlaH is considered essential for archaellal rotation [6] and these loci may therefore encode pilus-like structures rather than archaella. One Calditerrarchaeota genome (GCA_015520885.1) encoded two CRISPR-Cas systems (Type I-B and III-A) as well as a spacer-repeat set likely associated with the Type I-B system (Supplementary Table 5). Several other Calditerrarchaeota genomes (27%) encoded CRISPR-Cas spacer-repeat sets but did not possess identifiable CRISPR-Cas systems. Spacers from Calditerrarchaeota did not return high confidence hits against the NCBI_nr database.

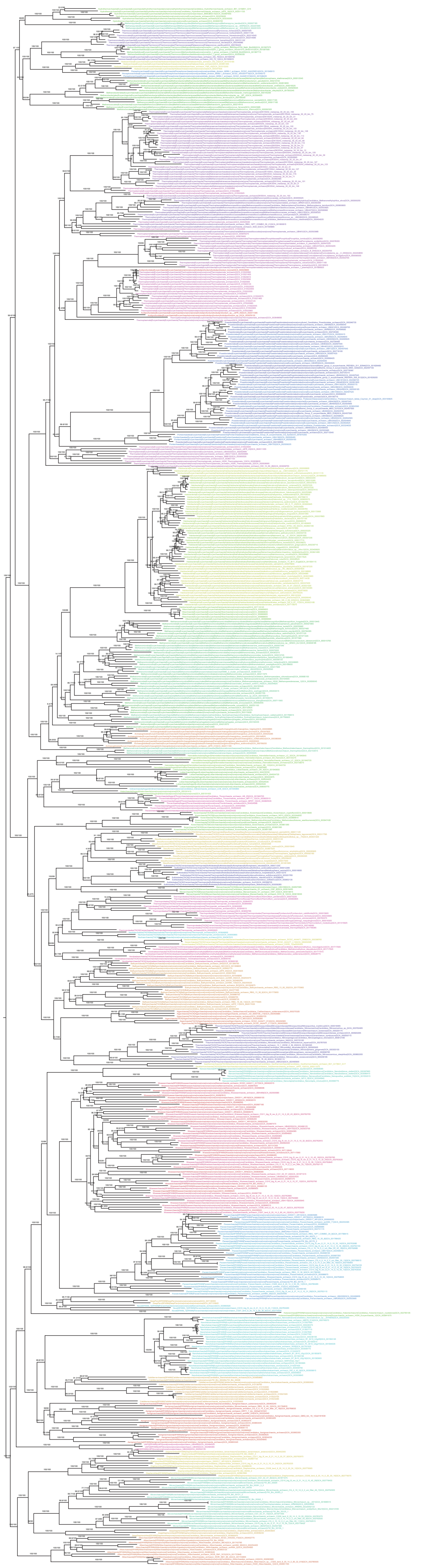

**Supplementary Figure 1 Phylogenetic placement of Calditerrarchaeota based on concatenated alignment of 50% top ranked marker genes and the 651 species set.** The alignment was trimmed with BMGE (Alignment length = 12,397 aa). A ML phylogenetic tree was inferred using IQ-Tree with the LG+C60+F+R model and ultrafast bootstrap approximation (left) and SH-like approximate likelihood tests (right), each run with 1000 replicates. The tree has been artificially rooted between DPANN and other Archaea. Scale bar: Average number of substitutions per site.

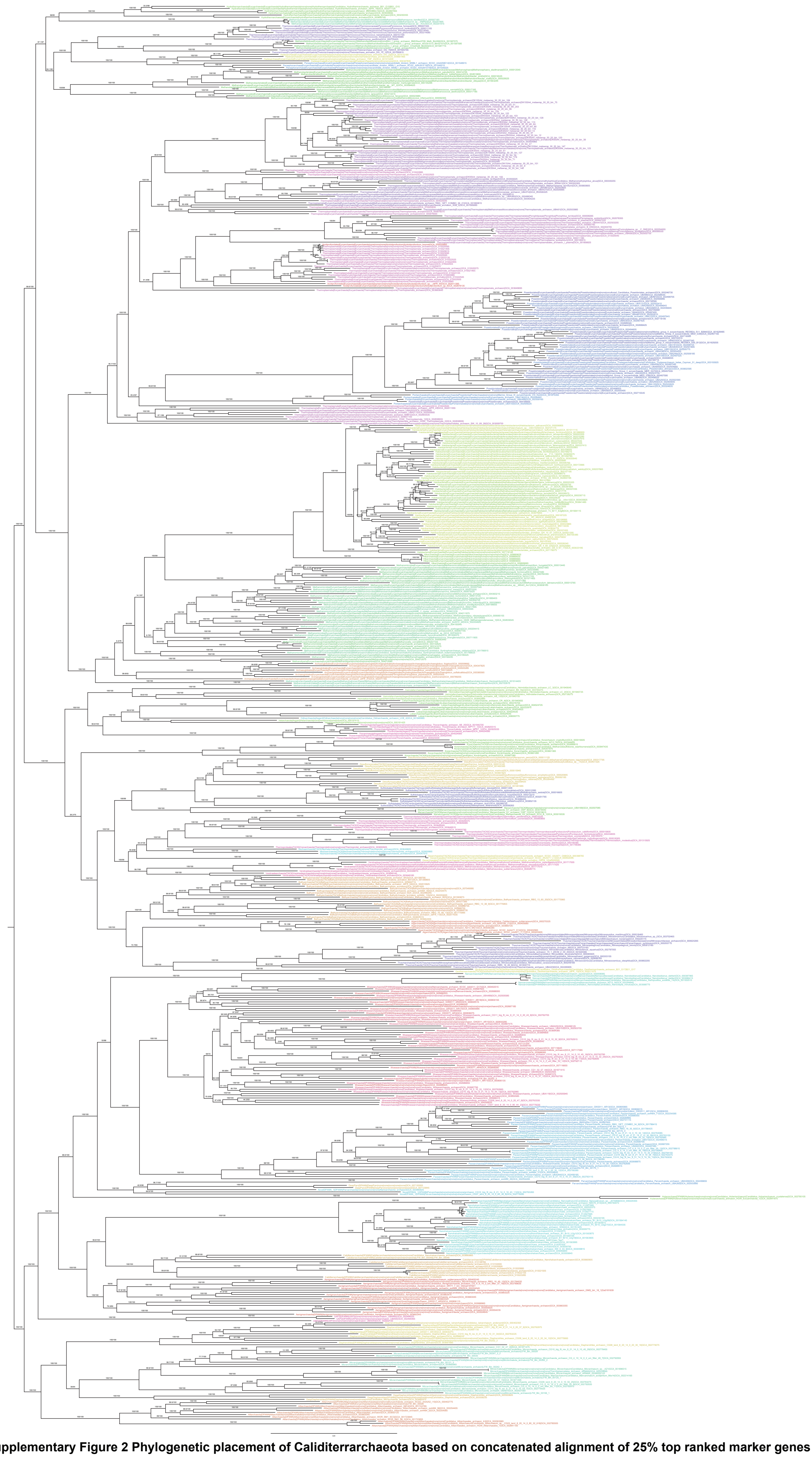

**Supplementary Figure 2 Phylogenetic placement of Caldarchaeales based on concatenated alignment of 25% top ranked marker genes and the 651 species set.** The alignment was trimmed with BMGE (Alignment length = 7,427 aa). A ML phylogenetic tree was inferred using IQ-Tree with the LG+C60+F+R model and ultrafast bootstrap approximation (left) and SH-like approximate likelihood tests (right), each run with 1000 replicates. The tree has been artificially rooted between DPANN and other Archaea. Scale bar: Average number of substitutions per site.

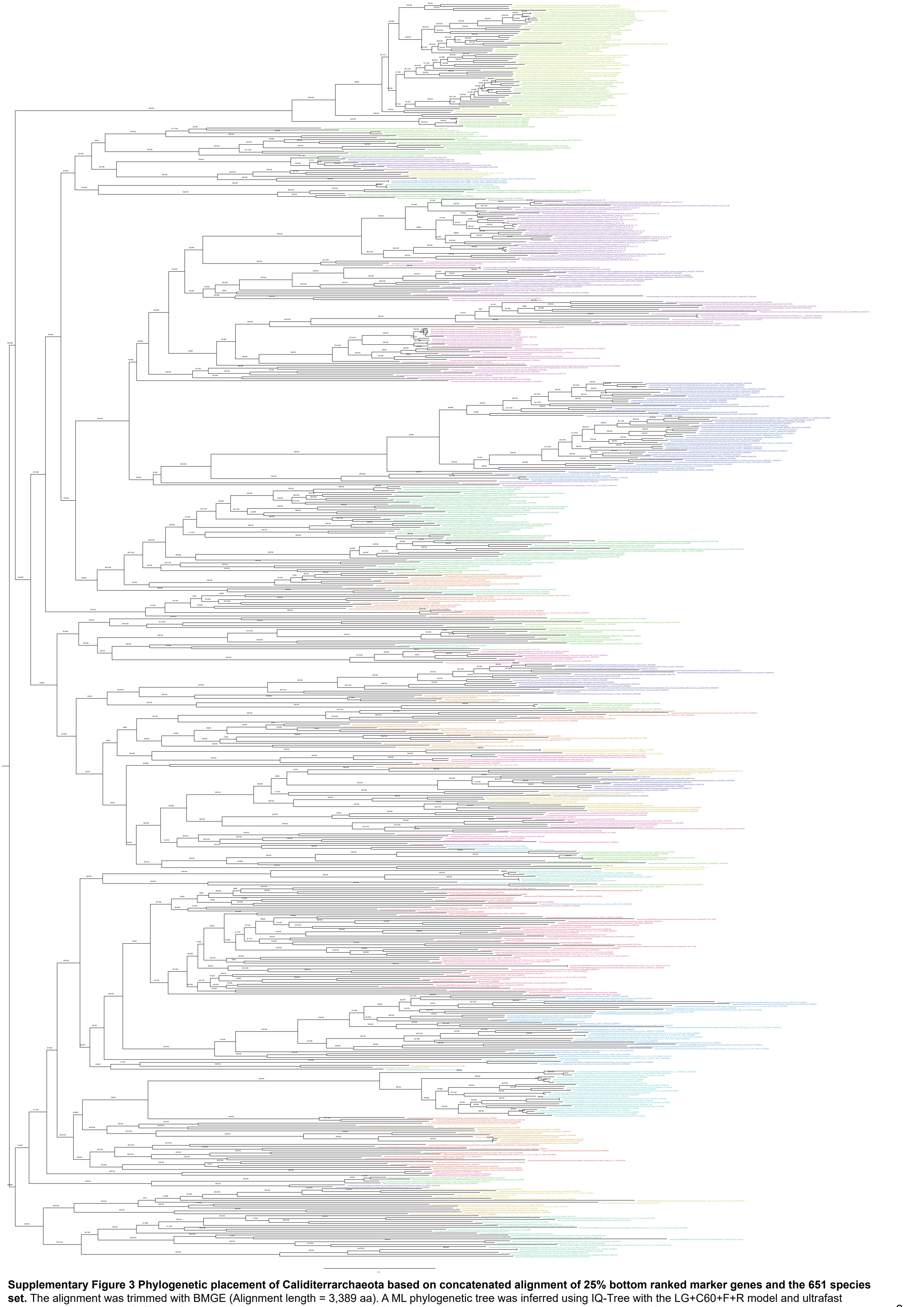

**Supplementary Figure 3 Phylogenetic placement of Caldititerrarchaeota based on concatenated alignment of 25% bottom ranked marker genes and the 651 species set.** The alignment was trimmed with BMGE (Alignment length = 3,389 aa). A ML phylogenetic tree was inferred using IQ-Tree with the LG+C60+F+R model and ultrafast bootstrap approximation (left) and SH-like approximate likelihood tests (right), each run with 1000 replicates. The tree has been artificially rooted between DPANN and other Archaea. Scale bar: Average number of substitutions per site.

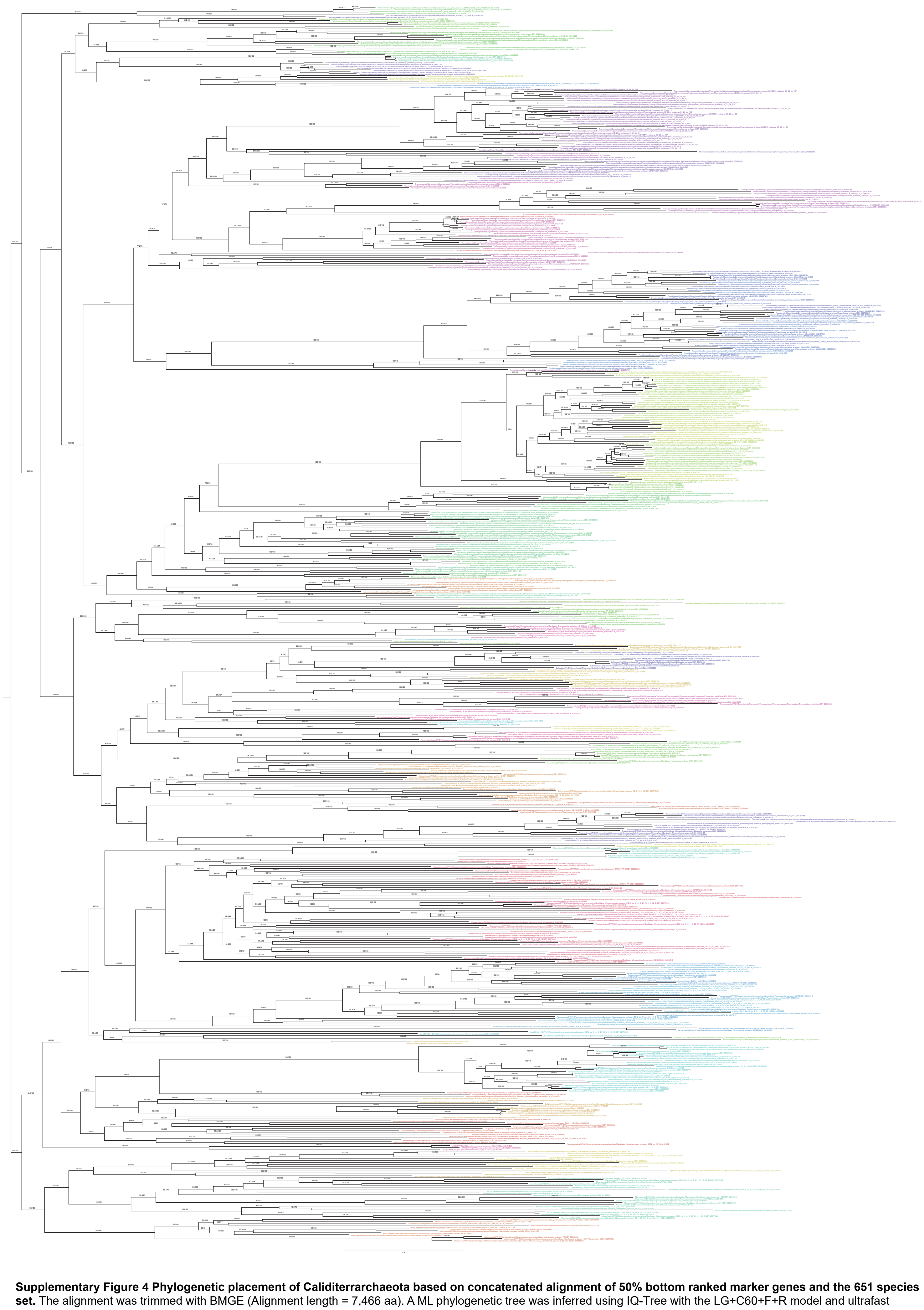

**Supplementary Figure 4 Phylogenetic placement of Calditerrarchaeota based on concatenated alignment of 50% bottom ranked marker genes and the 651 species set.** The alignment was trimmed with BMGE (Alignment length = 7,466 aa). A ML phylogenetic tree was inferred using IQ-Tree with the LG+C60+F+R model and ultrafast bootstrap approximation (left) and SH-like approximate likelihood tests (right), each run with 1000 replicates. The tree has been artificially rooted between DPANN and other Archaea. Scale bar: Average number of substitutions per site.

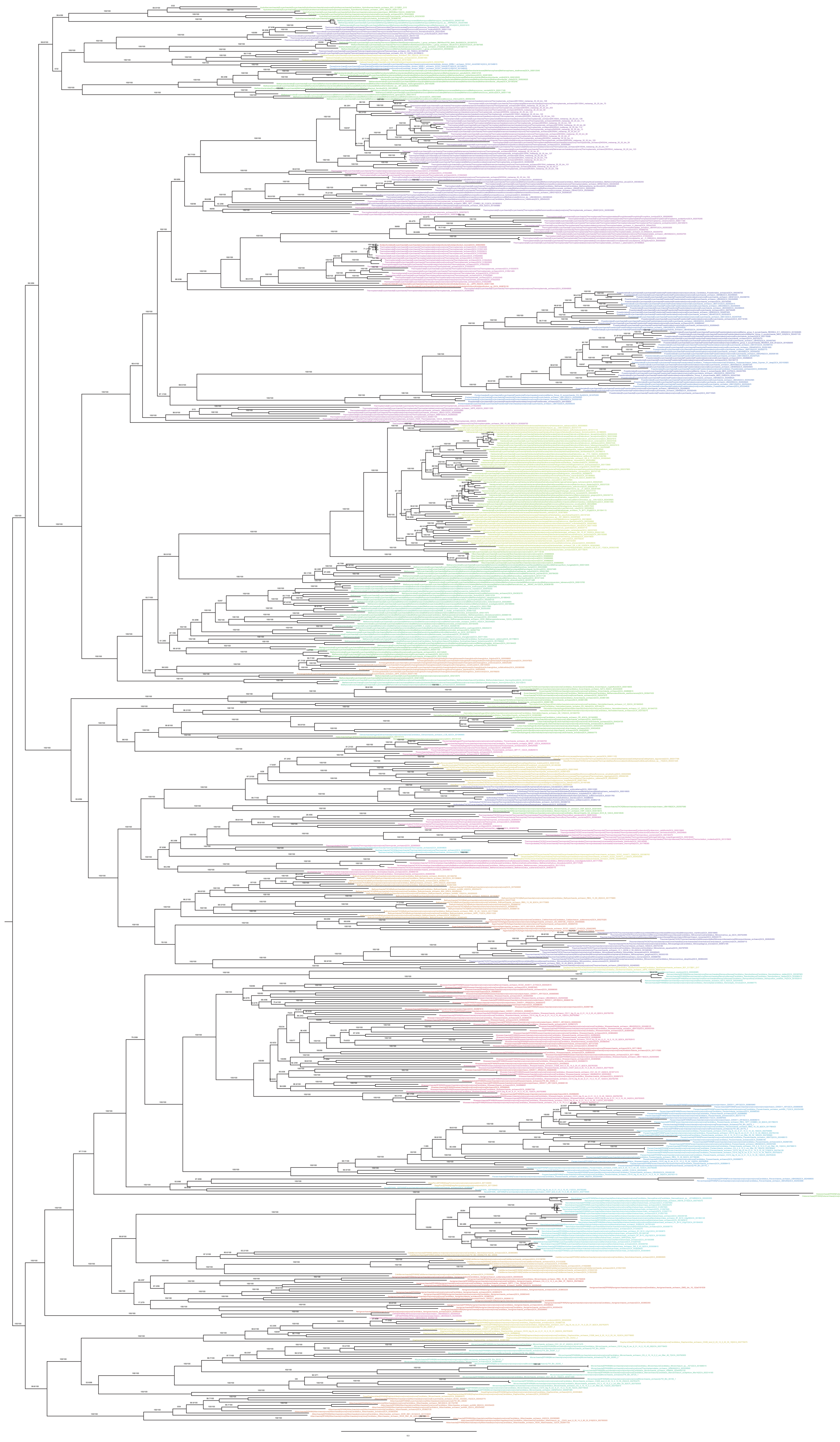

**Supplementary Figure 5 Phylogenetic placement of Calditerrarchaeota based on concatenated alignment of 50% top ranked marker genes and the 651 species set.** 10% of the fasted evolving sites were removed from the alignment with SlowFaster (Alignment length = 11,149 aa). A ML phylogenetic tree was inferred using IQ-Tree with the LG+C60+F+R model and ultrafast bootstrap approximation (left) and SH-like approximate likelihood tests (right), each run with 1000 replicates. The tree has been artificially rooted between DPANN and other Archaea. Scale bar: Average number of substitutions per site.

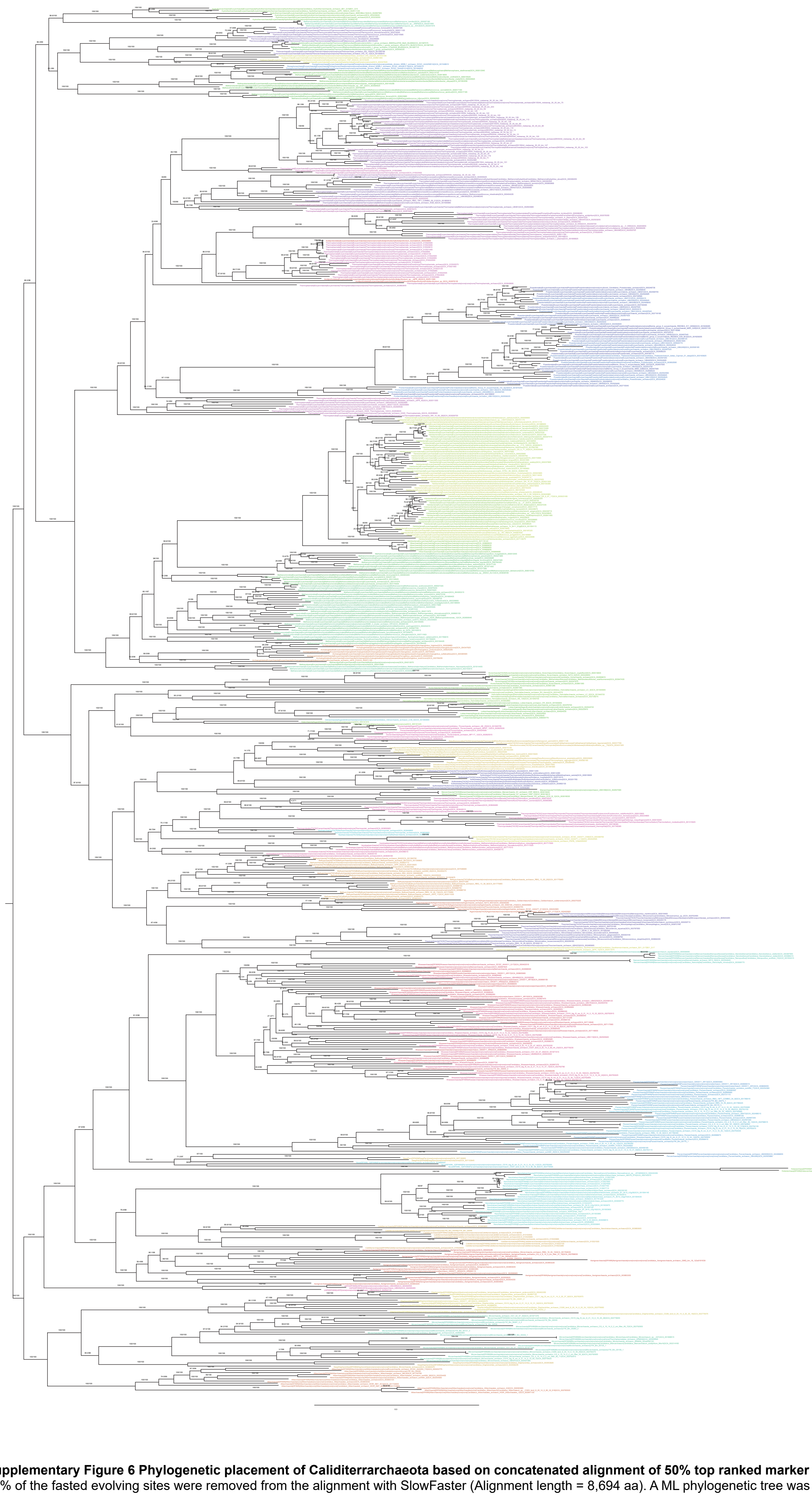

**Supplementary Figure 6 Phylogenetic placement of Calditerrarchaeota based on concatenated alignment of 50% top ranked marker genes and the 651 species set.** 30% of the fasted evolving sites were removed from the alignment with SlowFaster (Alignment length = 8,694 aa). A ML phylogenetic tree was inferred using IQ-Tree with the LG +C60+F+R model and ultrafast bootstrap approximation (left) and SH-like approximate likelihood tests (right), each run with 1000 replicates. The tree has been artificially rooted between DPANN and other Archaea. Scale bar: Average number of substitutions per site.

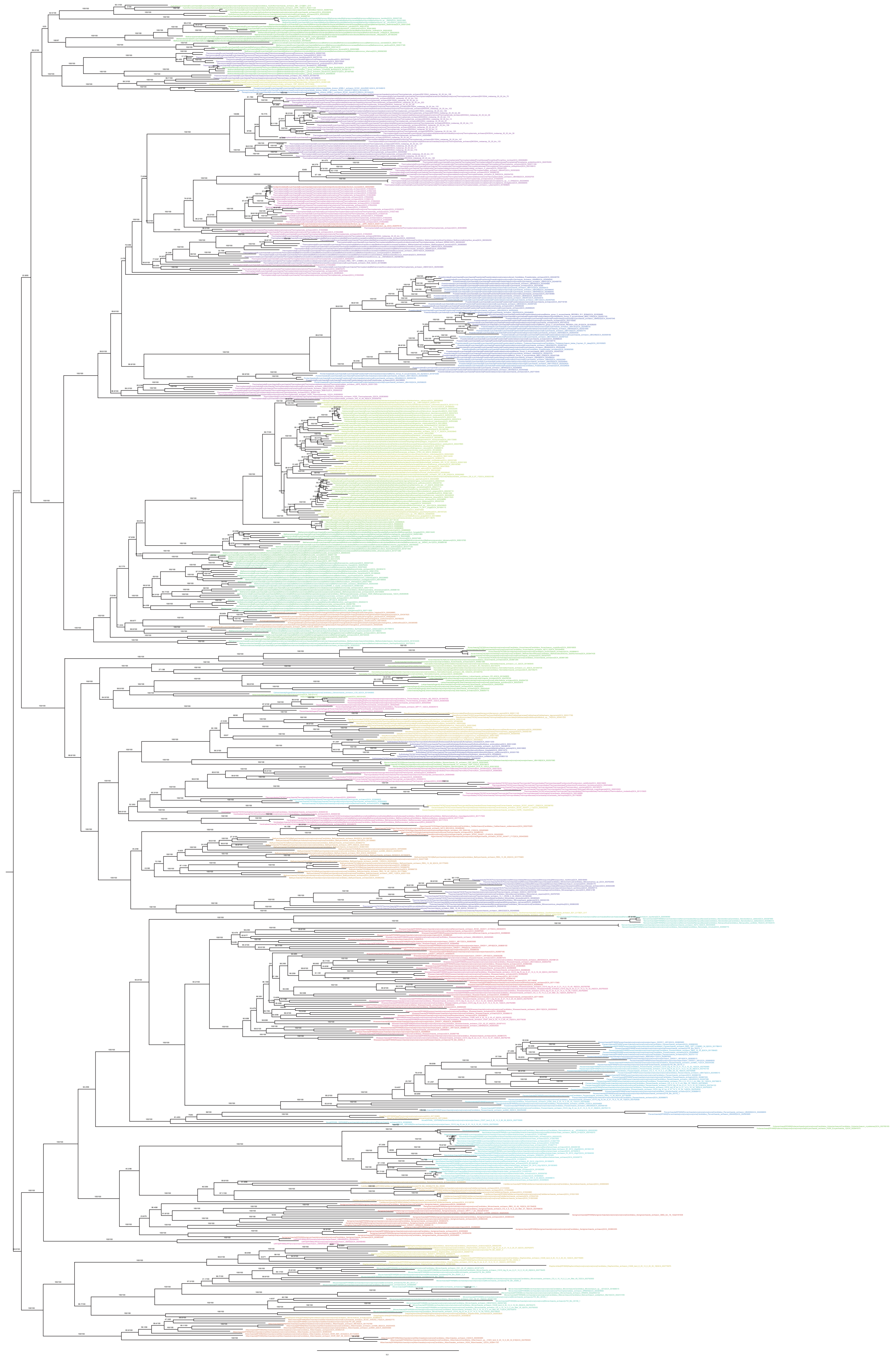

**Supplementary Figure 7 Phylogenetic placement of Caldíterrarchaeota based on concatenated alignment of 50% top ranked marker genes and the 651 species set.** 50% of the fasted evolving sites were removed from the alignment with SlowFaster (Alignment length = 6,169 aa). A ML phylogenetic tree was inferred using IQ-Tree with the LG+C60+F+R model and ultrafast bootstrap approximation (left) and SH-like approximate likelihood tests (right), each run with 1000 replicates. The tree has been artificially rooted between DPANN and other Archaea. Scale bar: Average number of substitutions per site.

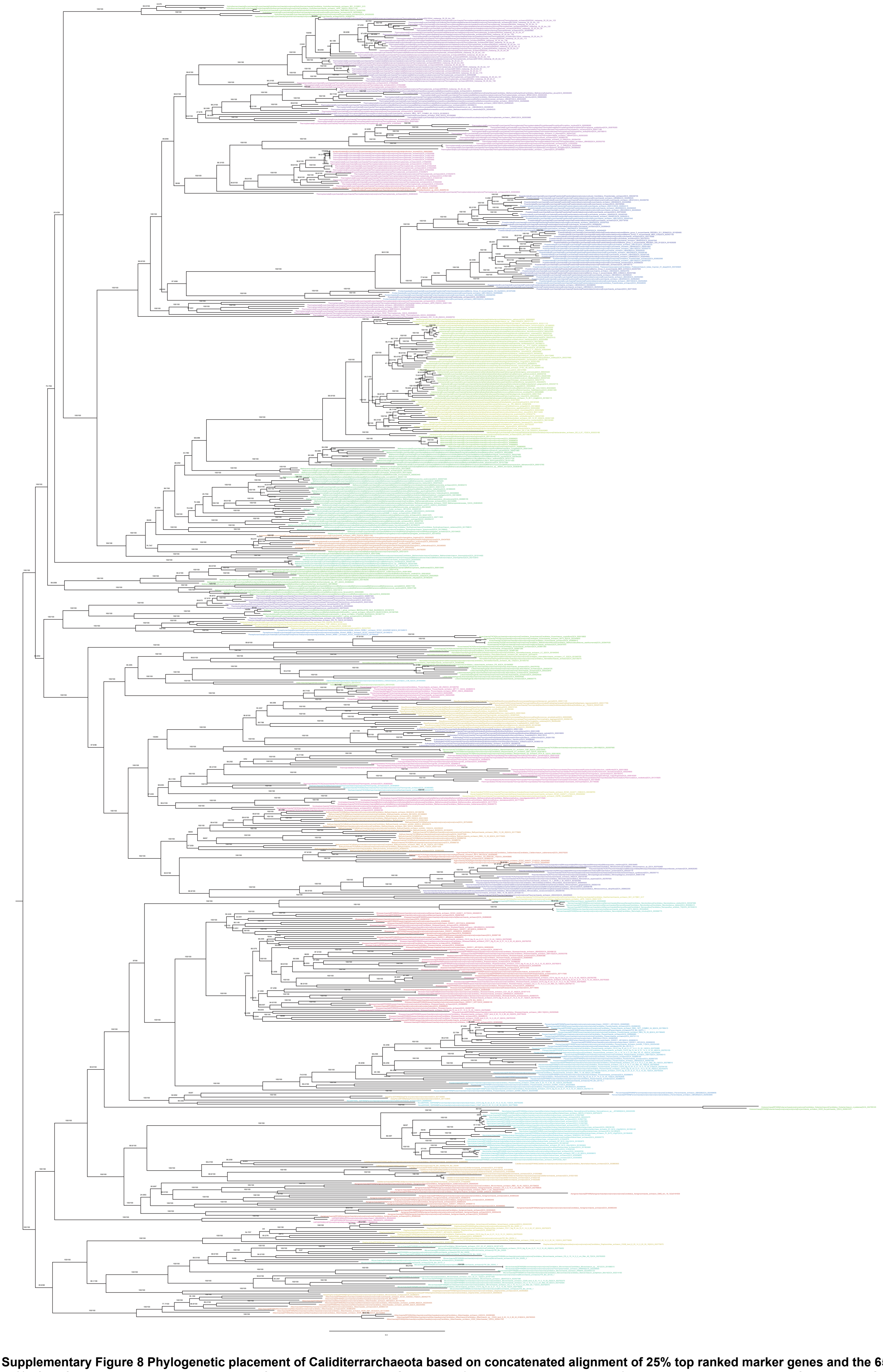

**Supplementary Figure 8 Phylogenetic placement of Calditerrarchaeota based on concatenated alignment of 25% top ranked marker genes and the 651 species set.** 10% of the fasted evolving sites were removed from the alignment with SlowFaster (Alignment length = 6,689 aa). A ML phylogenetic tree was inferred using IQ-Tree with the LG+C60+F+R model and ultrafast bootstrap approximation (left) and SH-like approximate likelihood tests (right), each run with 1000 replicates. The tree has been artificially rooted between DPANN and other Archaea. Scale bar: Average number of substitutions per site.

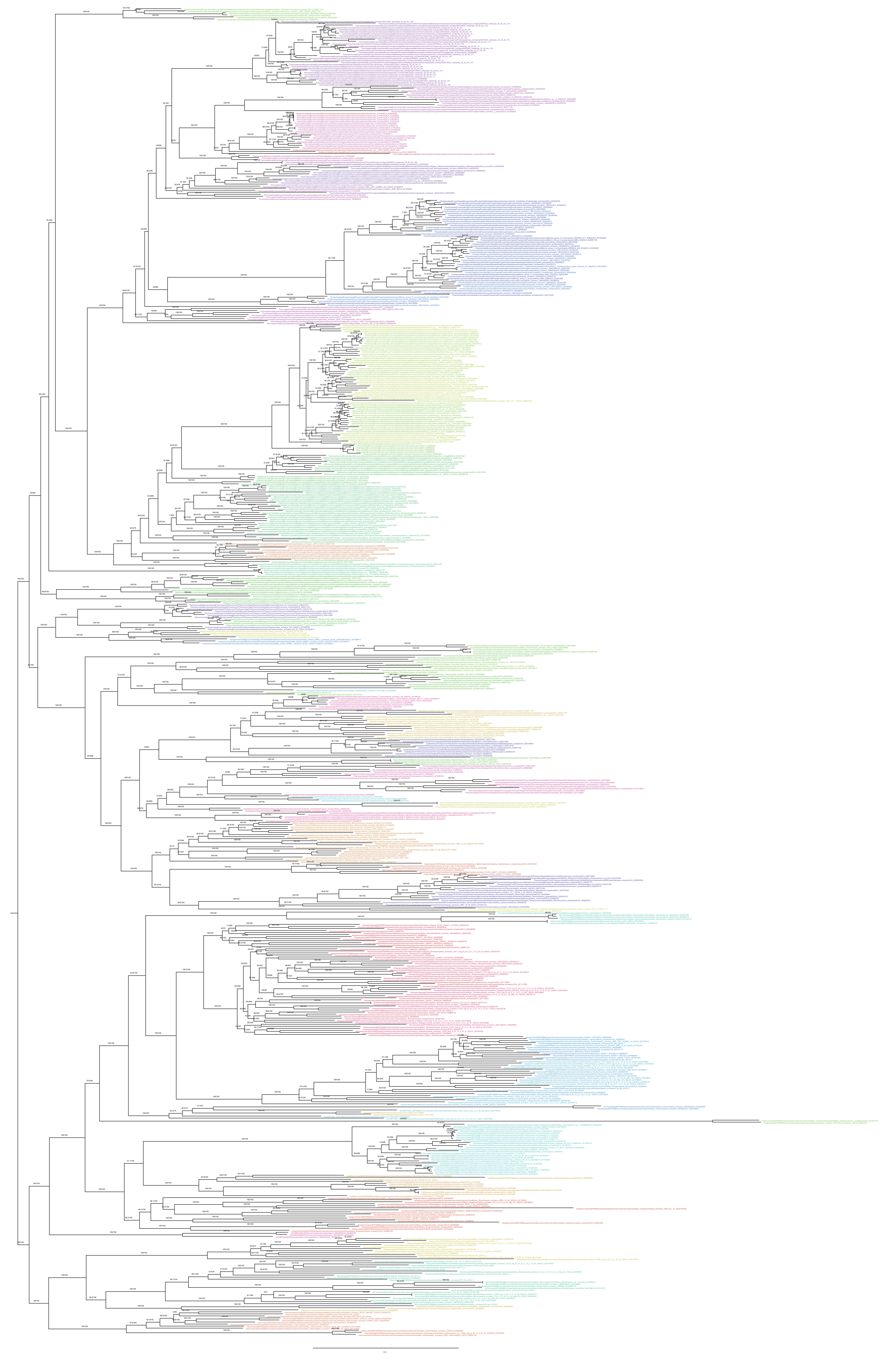

**Supplementary Figure 9 Phylogenetic placement of Calditerrarchaeota based on concatenated alignment of 25% top ranked marker genes and the 651 species set.** 30% of the fasted evolving sites were removed from the alignment with SlowFaster (Alignment length = 5,191 aa). A ML phylogenetic tree was inferred using IQ-Tree with the LG+C60+F+R model and ultrafast bootstrap approximation (left) and SH-like approximate likelihood tests (right), each run with 1000 replicates. The tree has been artificially rooted between DPANN and other Archaea. Scale bar: Average number of substitutions per site.

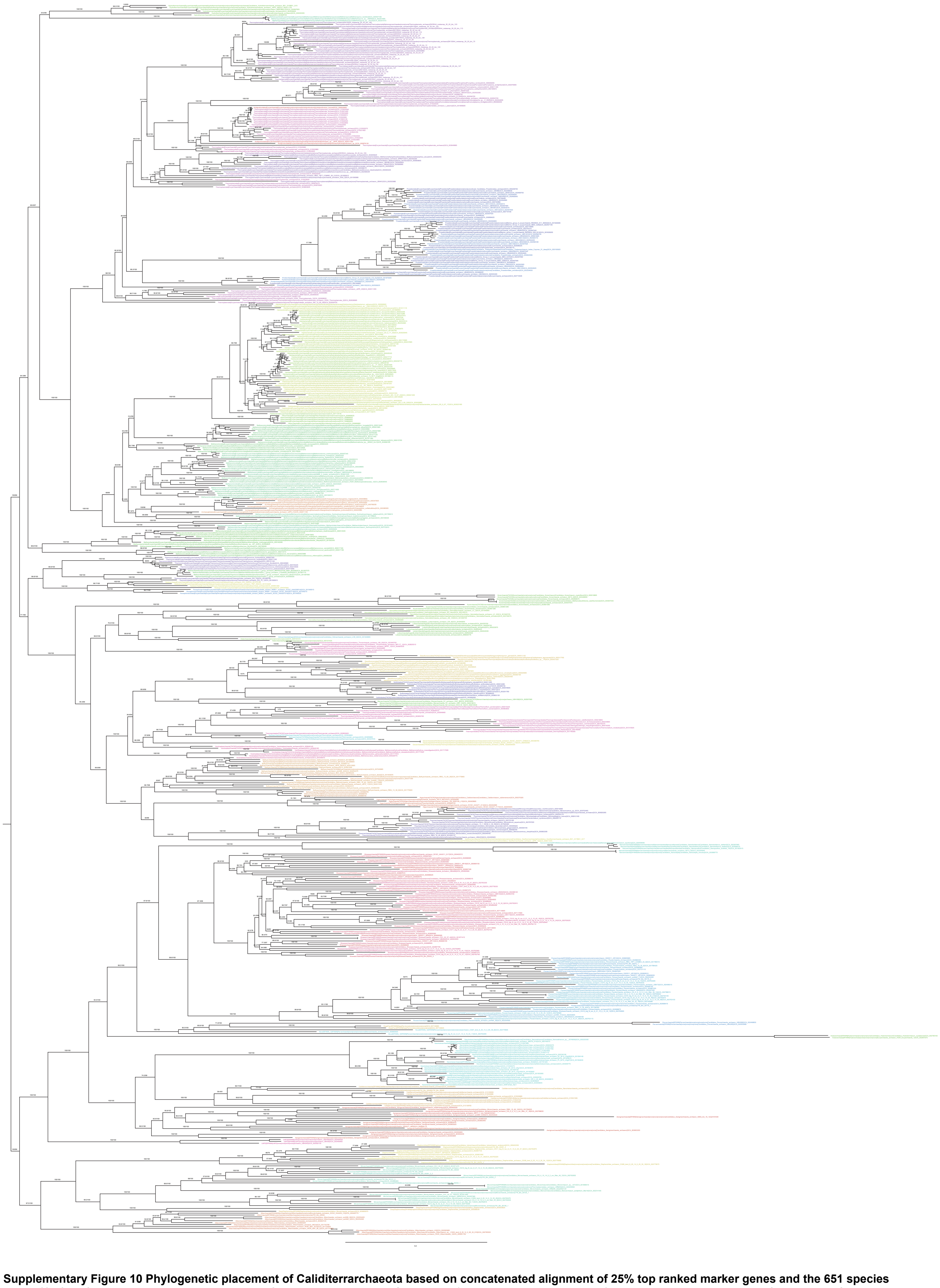

**Supplementary Figure 10 Phylogenetic placement of Calditerrarchaeota based on concatenated alignment of 25% top ranked marker genes and the 651 species set.** 50% of the fasted evolving sites were removed from the alignment with SlowFaster (Alignment length = 3,720 aa). A ML phylogenetic tree was inferred using IQ-Tree with the LG+C60+F+R model and ultrafast bootstrap approximation (left) and SH-like approximate likelihood tests (right), each run with 1000 replicates. The tree has been artificially rooted between DPANN and other Archaea. Scale bar: Average number of substitutions per site.

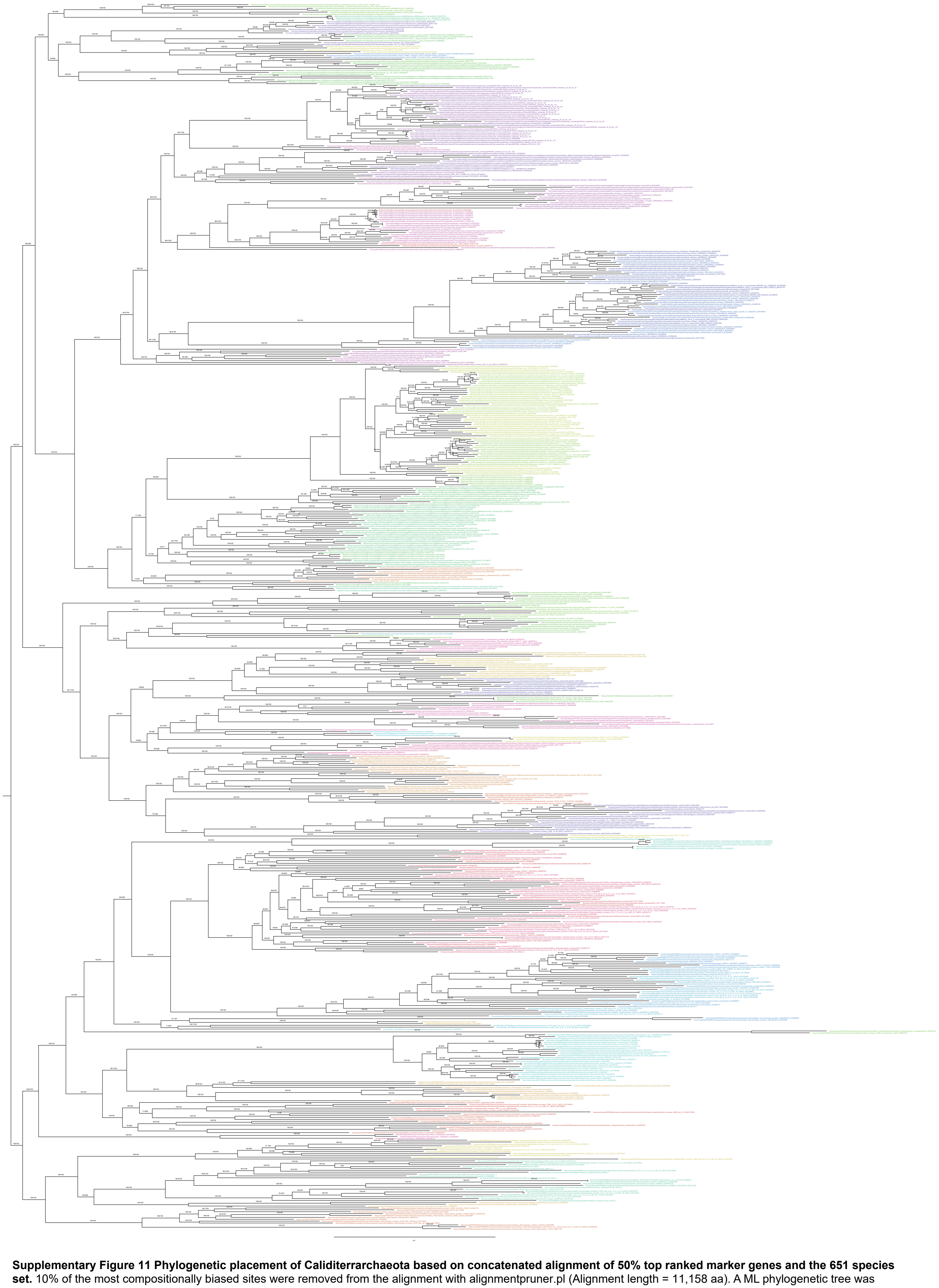

**Supplementary Figure 11 Phylogenetic placement of Calditerrarchaeota based on concatenated alignment of 50% top ranked marker genes and the 651 species set.** 10% of the most compositionally biased sites were removed from the alignment with alignmentpruner.pl (Alignment length = 11,158 aa). A ML phylogenetic tree was inferred using IQ-Tree with the LG+C60+F+R model and ultrafast bootstrap approximation (left) and SH-like approximate likelihood tests (right), each run with 1000 replicates. The tree has been artificially rooted between DPANN and other Archaea. Scale bar: Average number of substitutions per site.

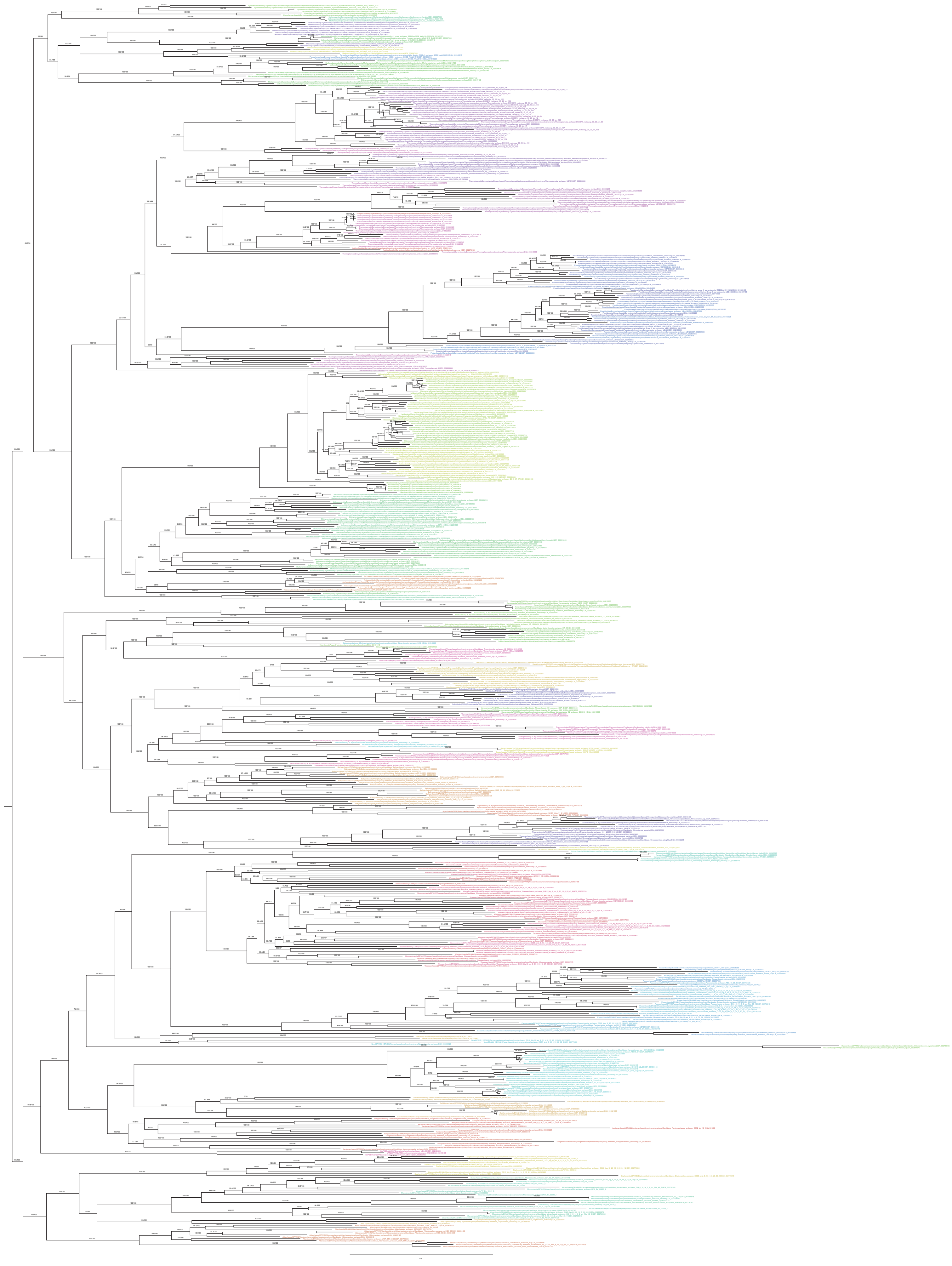

**Supplementary Figure 12 Phylogenetic placement of Calditerrarchaeota based on concatenated alignment of 30% top ranked marker genes and the 651 species set.** 30% of the most compositionally biased sites were removed from the alignment with alignmentpruner.pl (Alignment length = 8,678 aa). A ML phylogenetic tree was inferred using IQ-Tree with the LG+C60+F+R model and ultrafast bootstrap approximation (left) and SH-like approximate likelihood tests (right), each run with 1000 replicates. The tree has been artificially rooted between DPANN and other Archaea. Scale bar: Average number of substitutions per site.

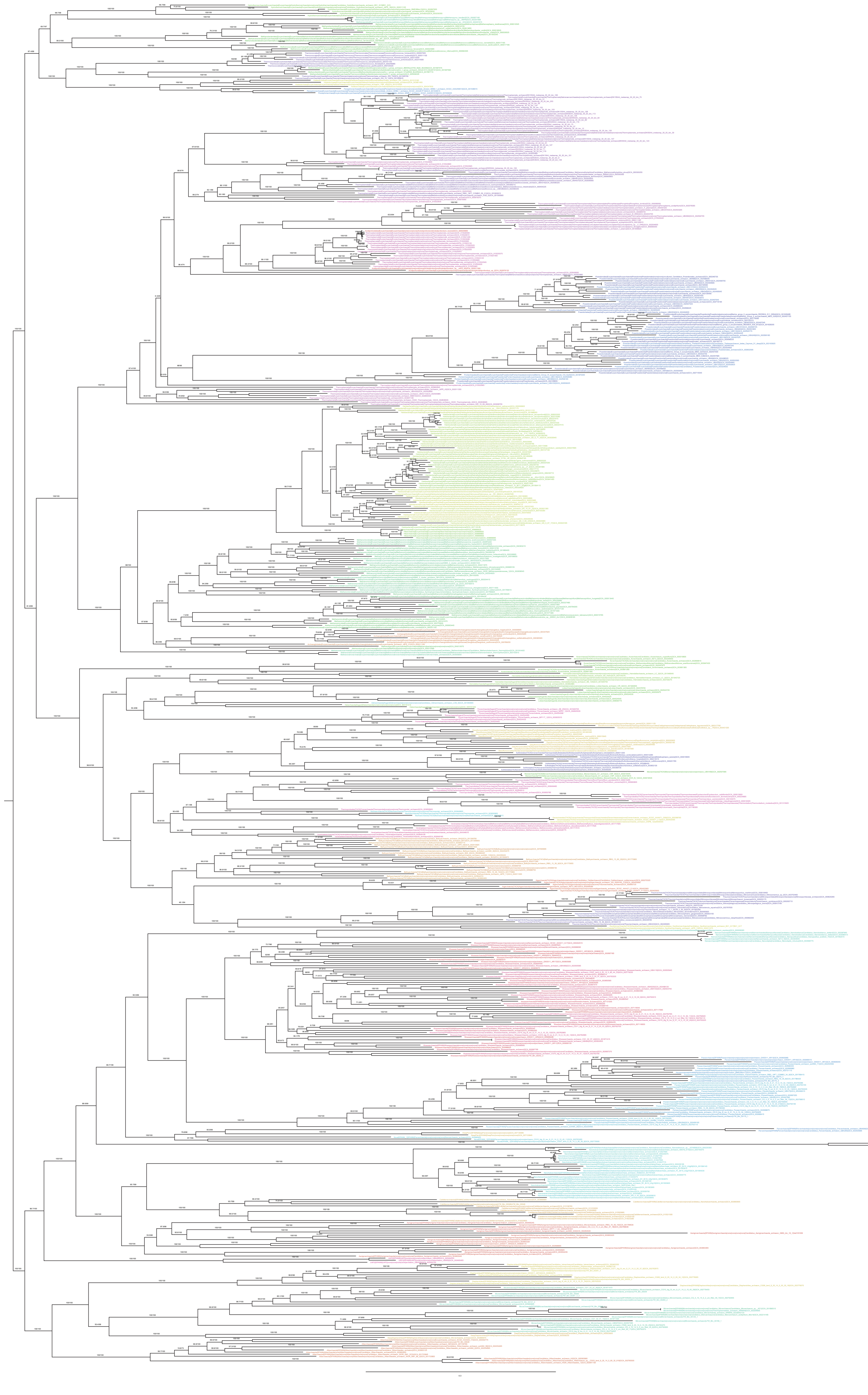

**Supplementary Figure 13 Phylogenetic placement of Caldiiterrarchaeota based on concatenated alignment of 50% top ranked marker genes and the 651 species set.** 50% of the most compositionally biased sites were removed from the alignment with alignmentpruner.pl (Alignment length = 6,199 aa). A ML phylogenetic tree was inferred using IQ-Tree with the LG+C60+F+R model and ultrafast bootstrap approximation (left) and SH-like approximate likelihood tests (right), each run with 1000 replicates. The tree has been artificially rooted between DPANN and other Archaea. Scale bar: Average number of substitutions per site.

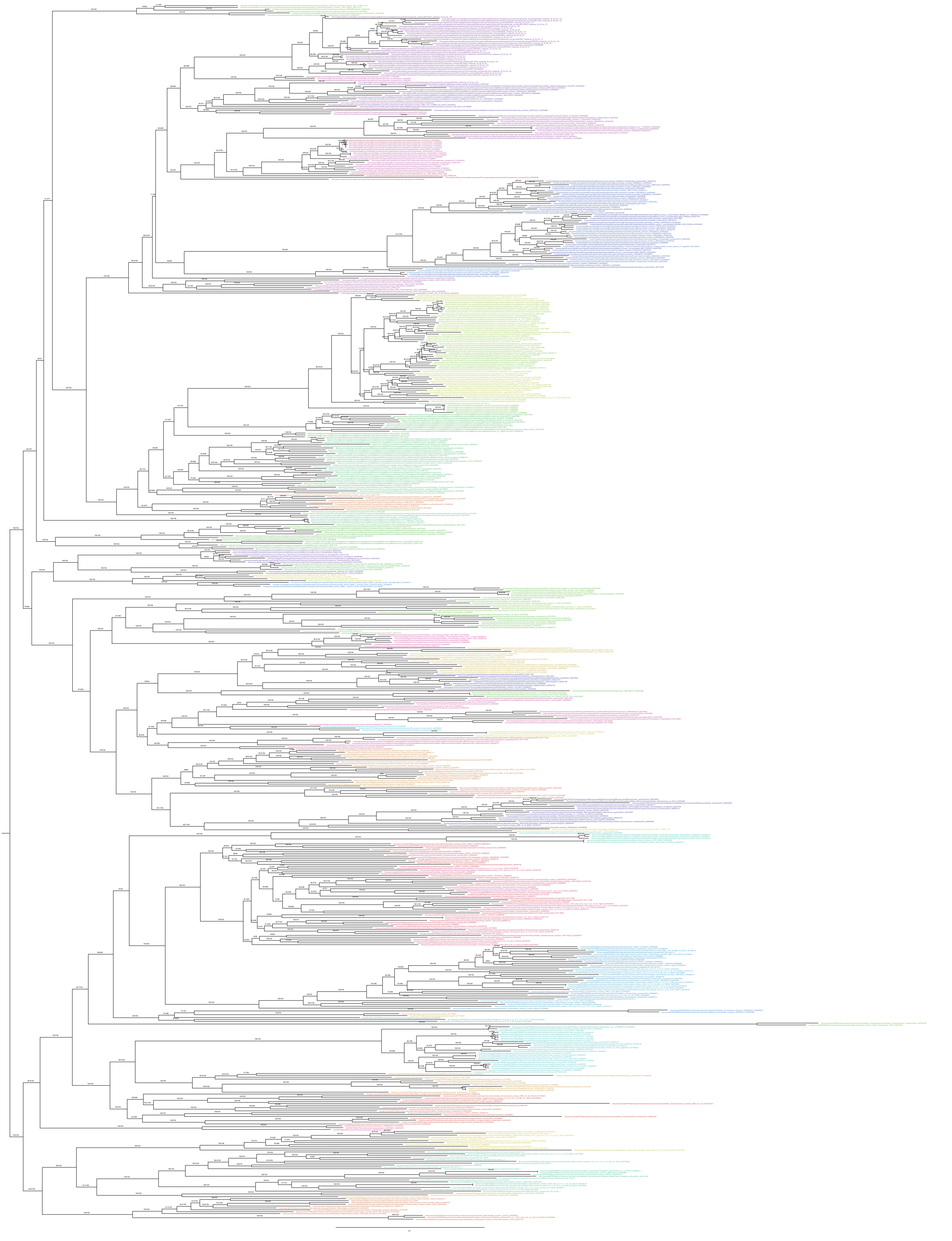

**Supplementary Figure 14 Phylogenetic placement of Calditerrarchaeota based on concatenated alignment of 25% top ranked marker genes and the 651 species set.** 10% of the most compositionally biased sites were removed from the alignment with alignmentpruner.pl (Alignment length = 6,685 aa). A ML phylogenetic tree was inferred using IQ-Tree with the LG+C60+F+R model and ultrafast bootstrap approximation (left) and SH-like approximate likelihood tests (right), each run with 1000 replicates. The tree has been artificially rooted between DPANN and other Archaea. Scale bar: Average number of substitutions per site.

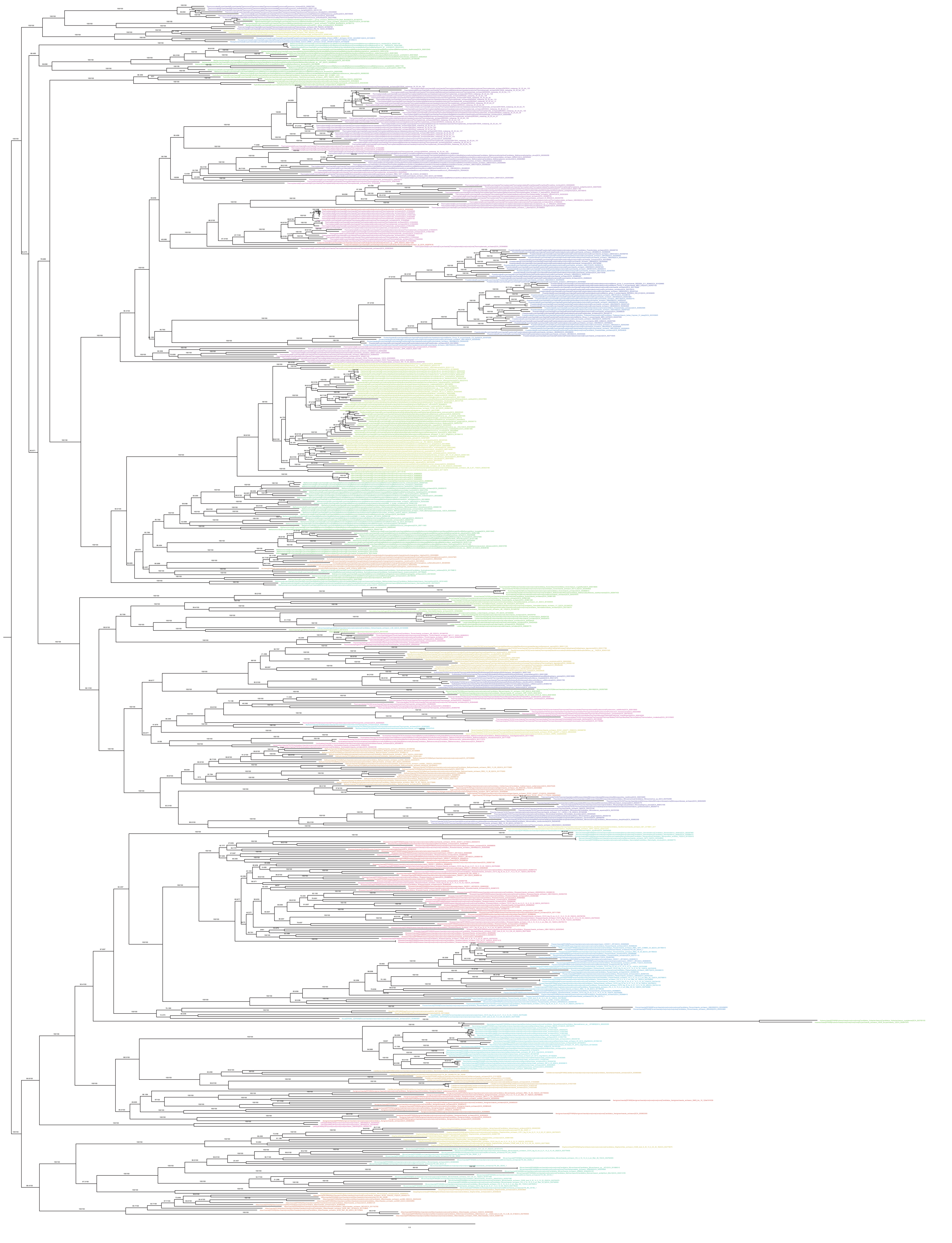

**Supplementary Figure 15 Phylogenetic placement of Calditerrarchaeota based on concatenated alignment of 25% top ranked marker genes and the 651 species set.** 30% of the most compositionally biased sites were removed from the alignment with alignmentpruner.pl (Alignment length = 5,199 aa). A ML phylogenetic tree was inferred using IQ-Tree with the LG+C60+F+R model and ultrafast bootstrap approximation (left) and SH-like approximate likelihood tests (right), each run with 1000 replicates. The tree has been artificially rooted between DPANN and other Archaea. Scale bar: Average number of substitutions per site.

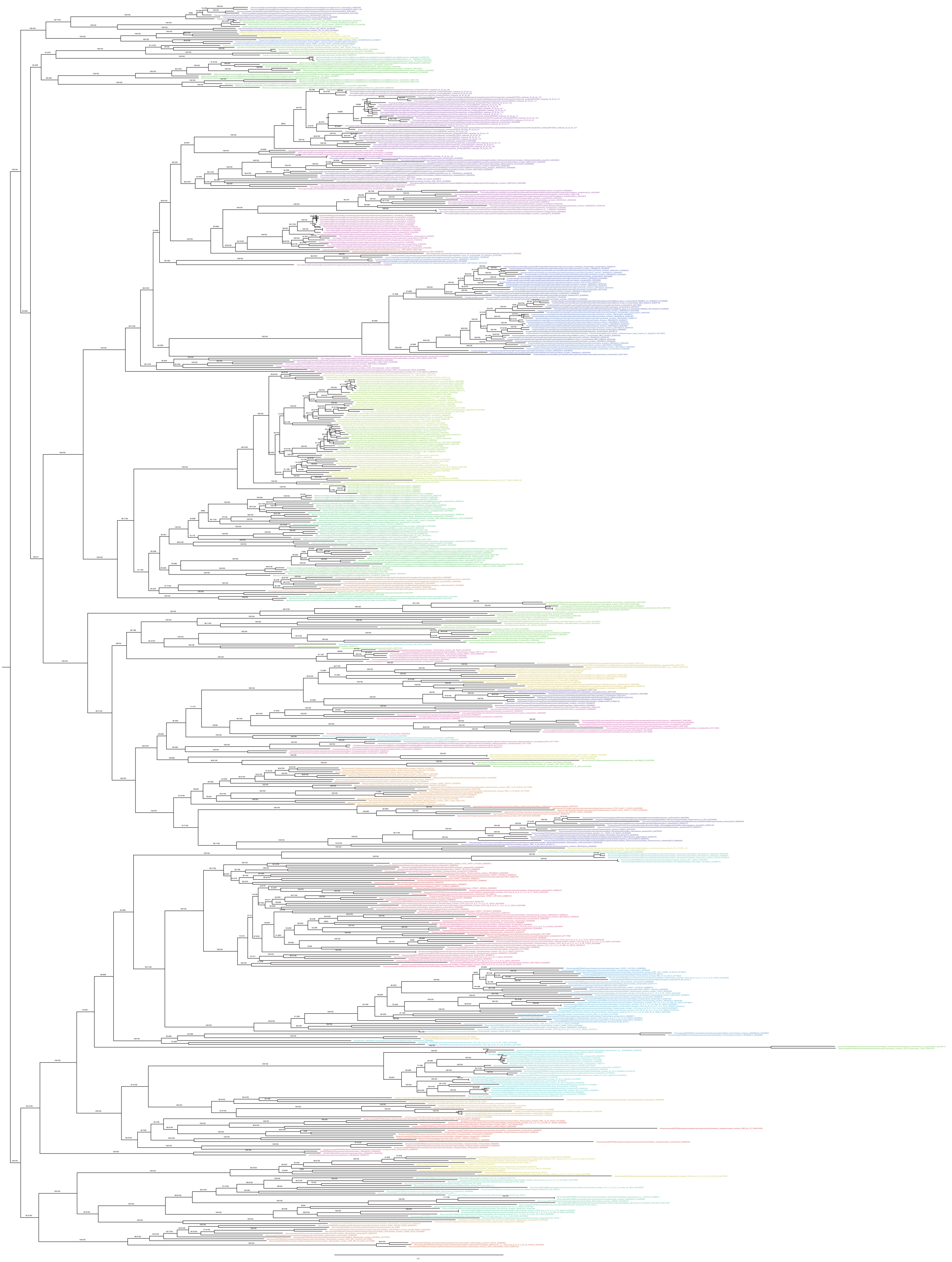

**Supplementary Figure 16 Phylogenetic placement of Calditerrarchaeota based on concatenated alignment of 25% top ranked marker genes and the 651 species set.** 50% of the most compositionally biased sites were removed from the alignment with alignmentpruner.pl (Alignment length = 3,714 aa). A ML phylogenetic tree was inferred using IQ-Tree with the LG+C60+F+R model and ultrafast bootstrap approximation (left) and SH-like approximate likelihood tests (right), each run with 1000 replicates. The tree has been artificially rooted between DPANN and other Archaea. Scale bar: Average number of substitutions per site.

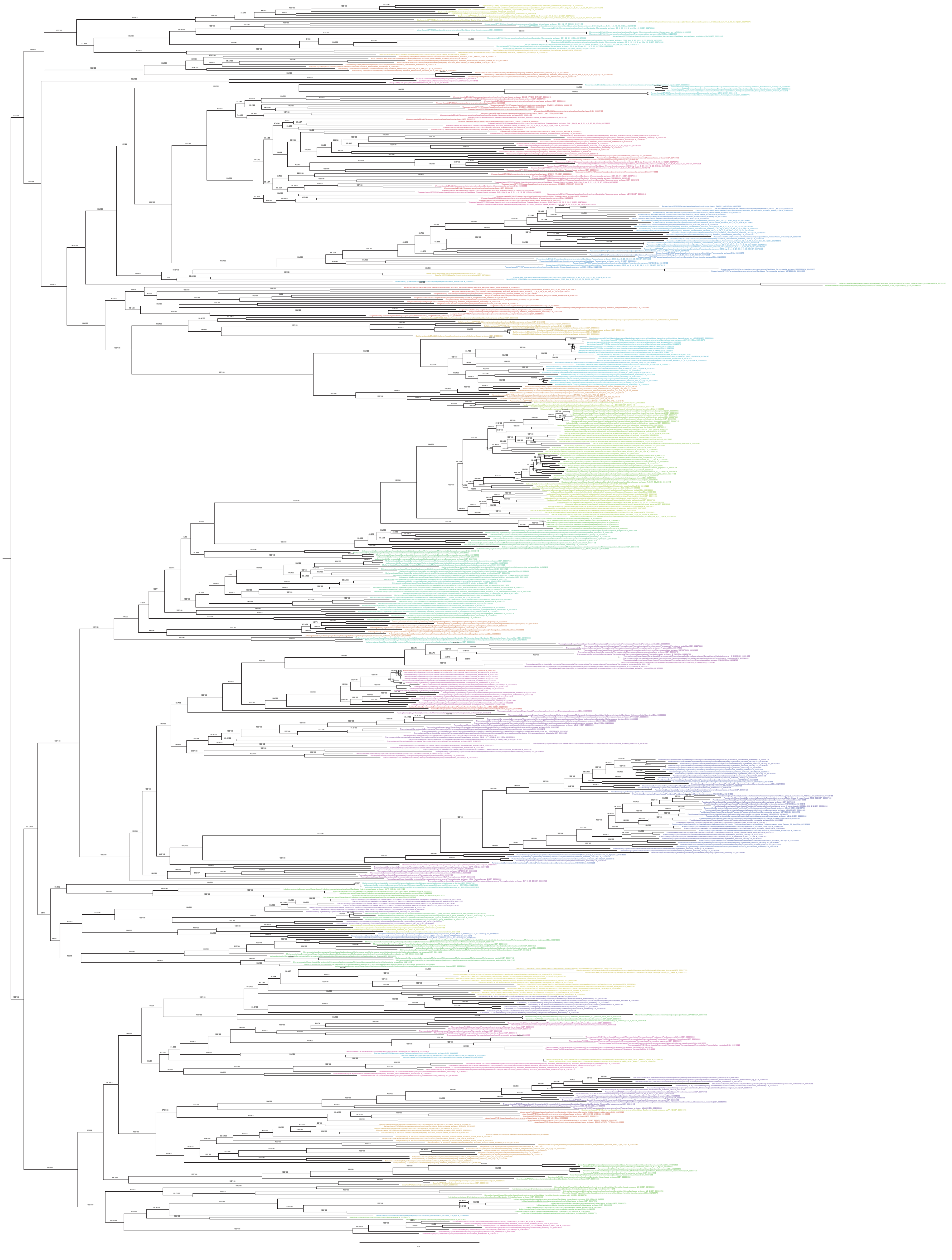

**Supplementary Figure 17 Phylogenetic placement of Calditerrarchaeota based on concatenated alignment of 50% top ranked marker genes and the 659 species set (Main set with added *Asbonarchaeaceae*).** The alignment was trimmed with BMGE (Alignment length = 12,432 aa). A ML phylogenetic tree was inferred using IQ-Tree with the LG+C60+F+R model and ultrafast bootstrap approximation (left) and SH-like approximate likelihood tests (right), each run with 1000 replicates. The tree has been artificially rooted between DPANN and other Archaea. Scale bar: Average number of substitutions per site.

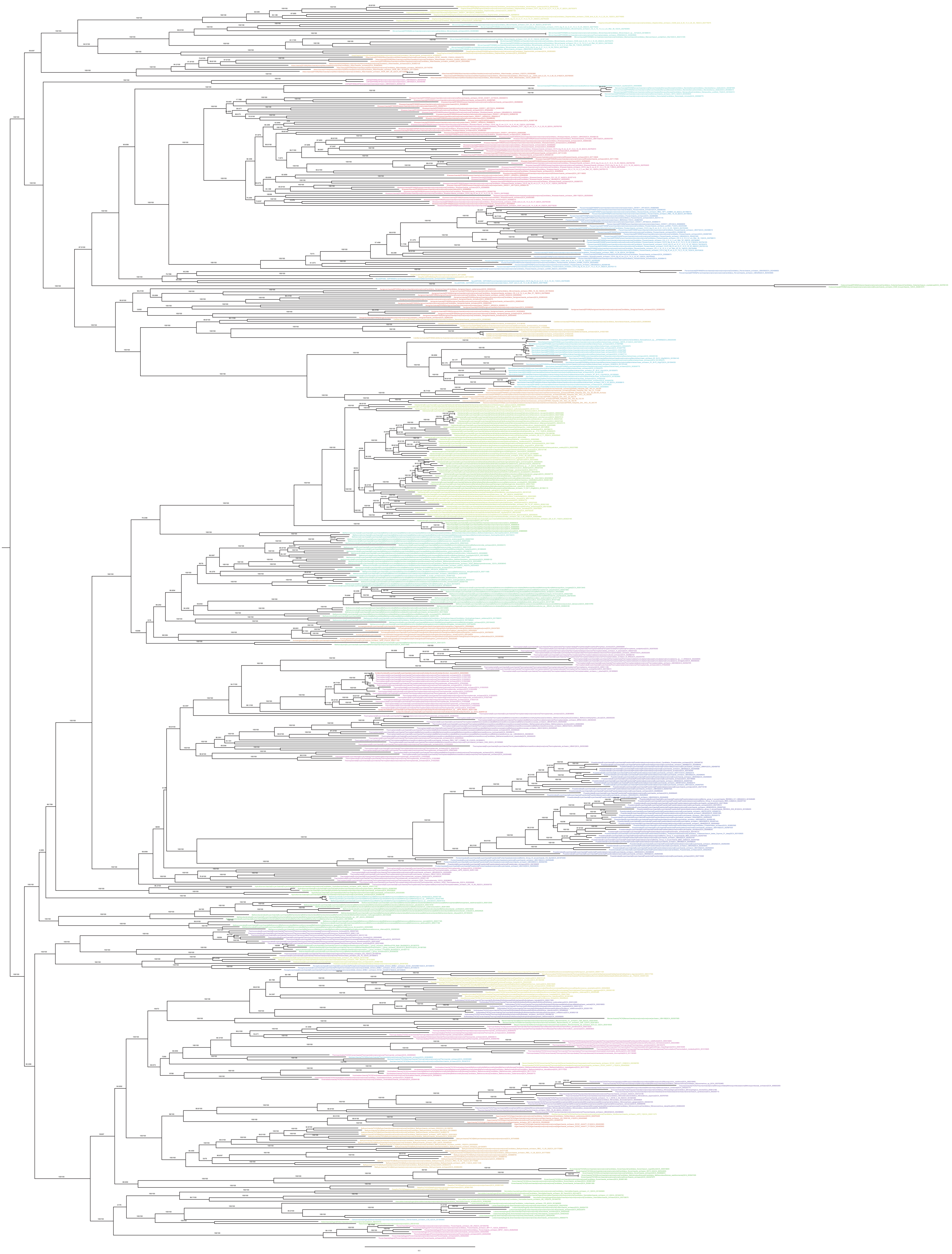

**Supplementary Figure 18 Phylogenetic placement of Calditerrarchaeota based on concatenated alignment of 25% top ranked marker genes and the 659 species set (Main set with added *Asbonarchaeaceae*).** The alignment was trimmed with BMGE (Alignment length = 7,473 aa). A ML phylogenetic tree was inferred using IQ-Tree with the LG+C60+F+R model and ultrafast bootstrap approximation (left) and SH-like approximate likelihood tests (right), each run with 1000 replicates. The tree has been artificially rooted between DPANN and other Archaea. Scale bar: Average number of substitutions per site.

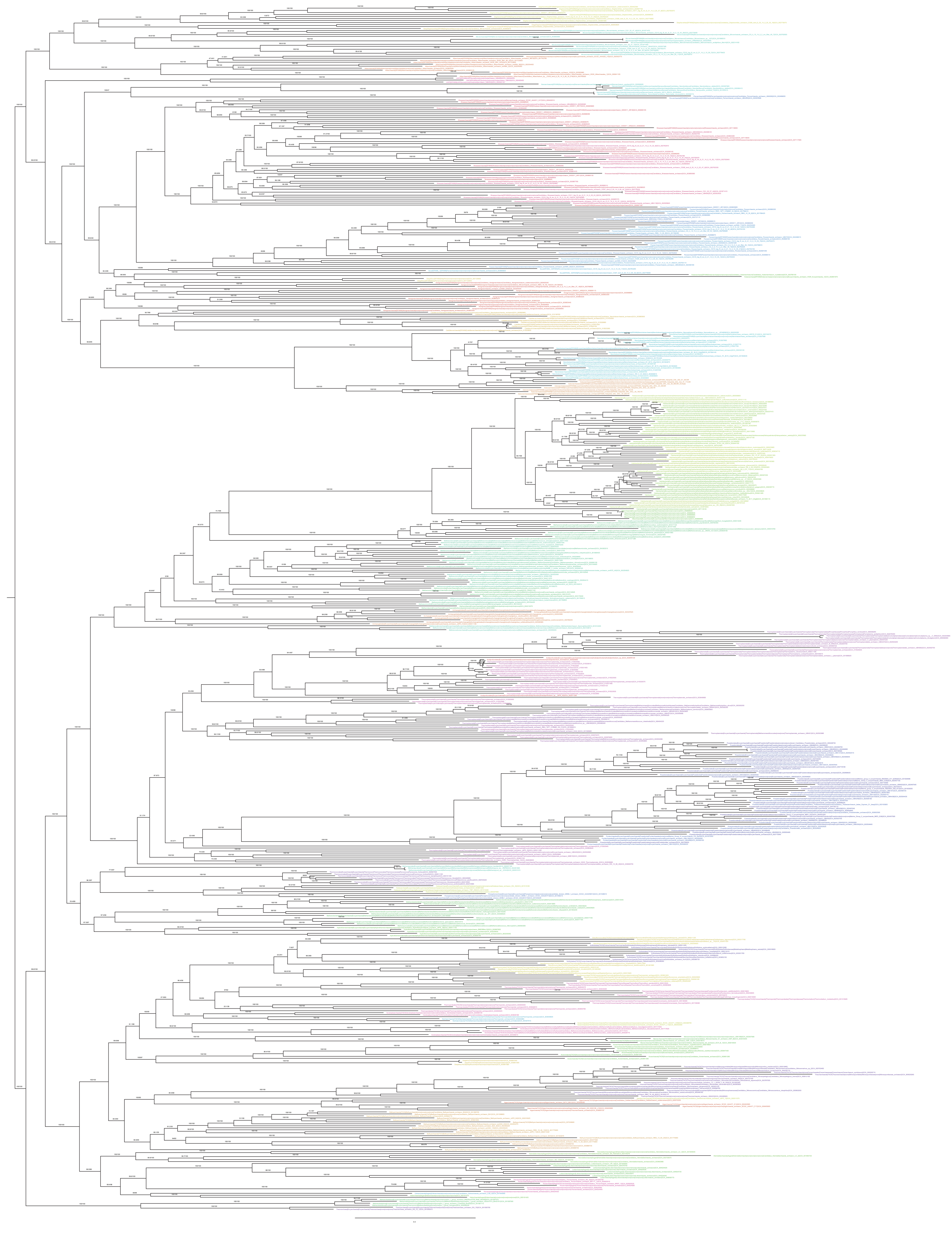

**Supplementary Figure 19 Phylogenetic placement of Calditerrarchaeota based on concatenated alignment of 50% bottom ranked marker genes and the 659 species set (Main set with added *Asbonarchaeaceae*).** The alignment was trimmed with BMGE (Alignment length = 7,569 aa). A ML phylogenetic tree was inferred using IQ-Tree with the LG+C60+F+R model and ultrafast bootstrap approximation (left) and SH-like approximate likelihood tests (right), each run with 1000 replicates. The tree has been artificially rooted between DPANN and other Archaea. Scale bar: Average number of substitutions per site.

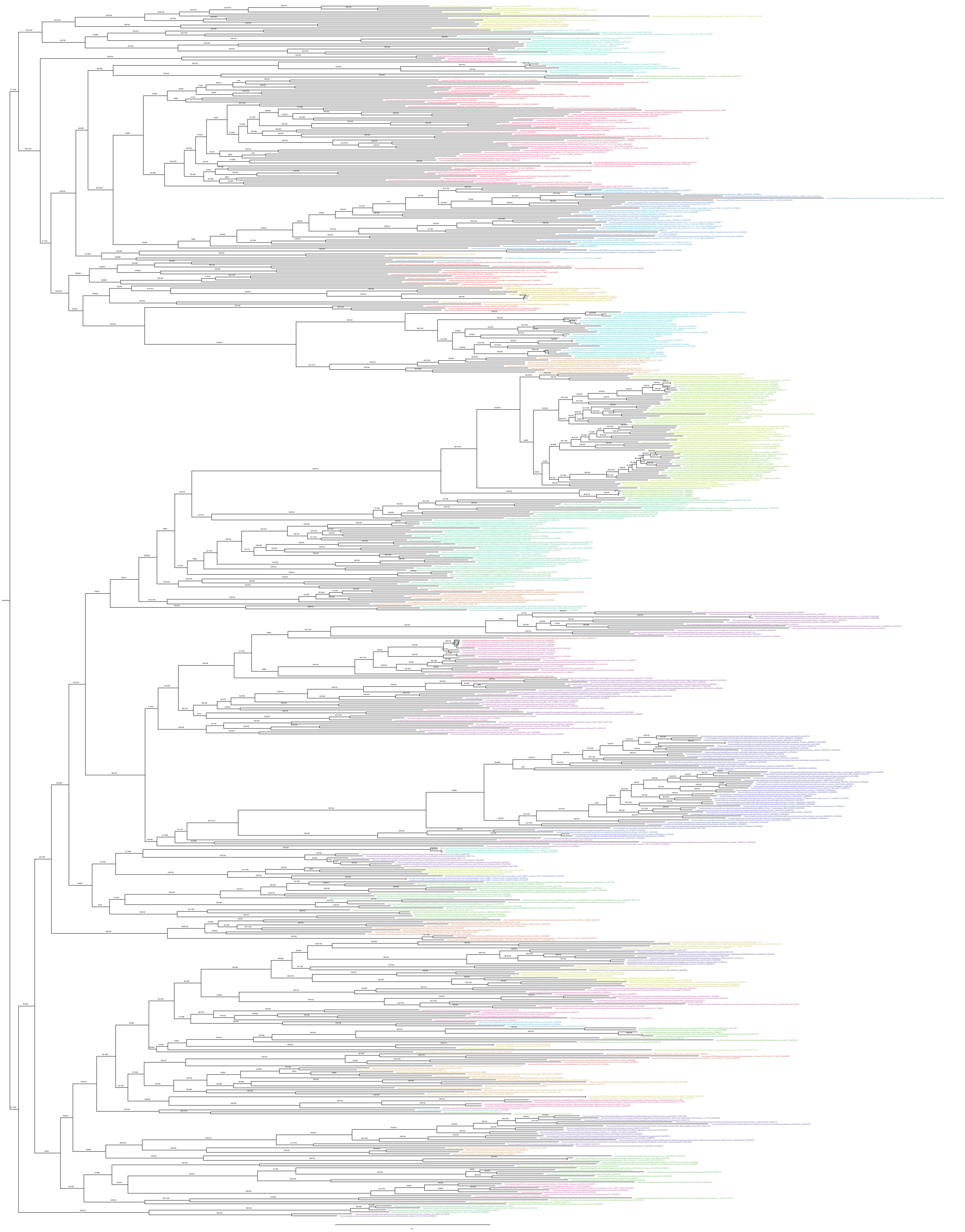

**Supplementary Figure 20 Phylogenetic placement of Calditerrarchaeota based on concatenated alignment of 25% bottom ranked marker genes and the 659 species set (Main set with added *Asbonarchaeaceae*).** The alignment was trimmed with BMGE (Alignment length = 3,437 aa). A ML phylogenetic tree was inferred using IQ-Tree with the LG+C60+F+R model and ultrafast bootstrap approximation (left) and SH-like approximate likelihood tests (right), each run with 1000 replicates. The tree has been artificially rooted between DPANN and other Archaea. Scale bar: Average number of substitutions per site.

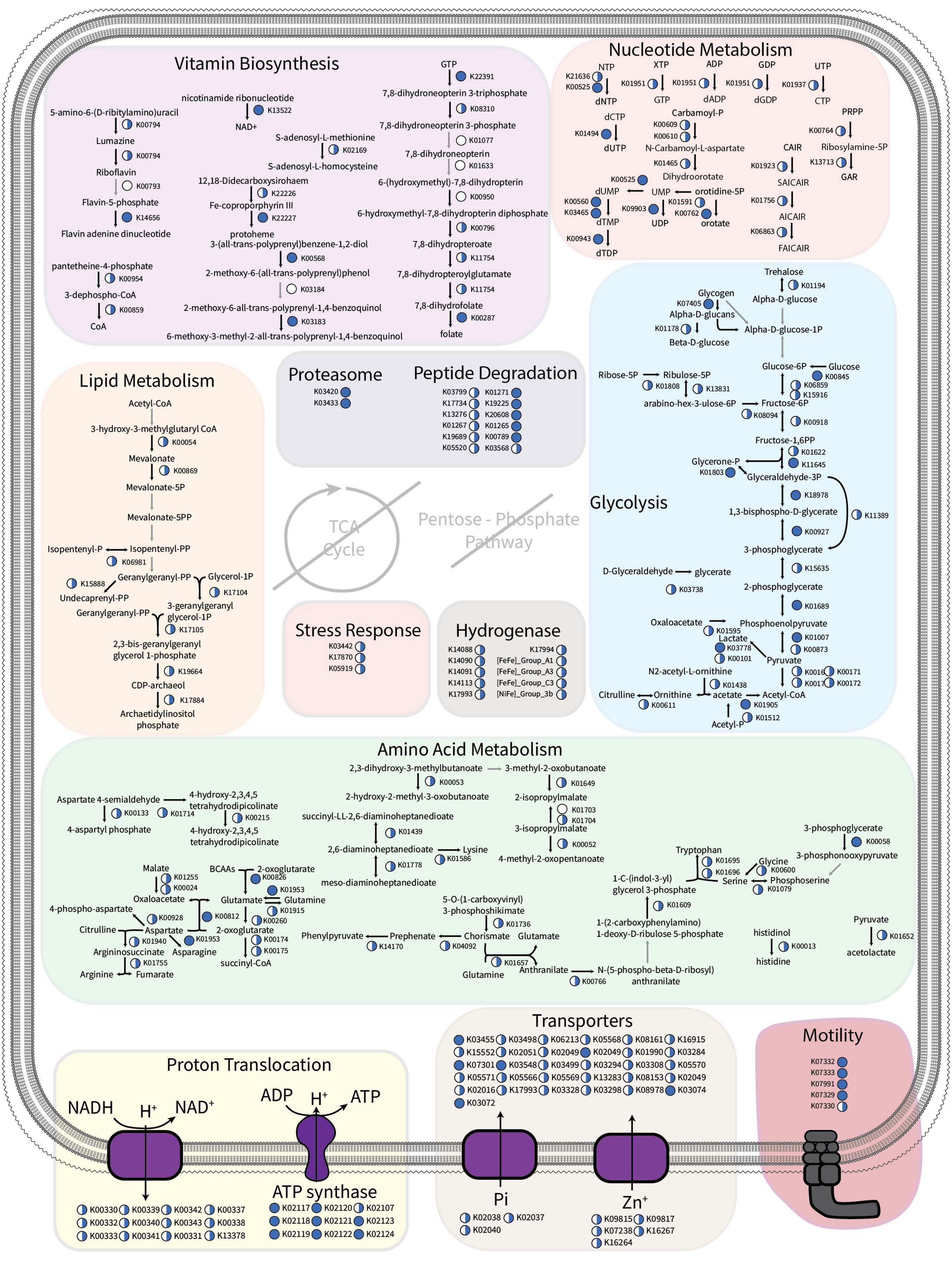


**Supplementary Figure 21 Genome content of Calditerrarchaeota MAGs.** Full circles indicate gene presence in >=50% Calditerrarchaeota MAGs. Half circles indicate presence in at least one but <50% of Calditerrarchaeota MAGs. Greyed out pathways indicate pathways or genes completely absent from Calditerrarchaeota MAGs included in the dataset. The full annotations for Calditerrarchaeota MAGs analysed in this study are available in Supplementary Table 3 and the specific genes presented in this figure are summarised in Supplementary Table 4.

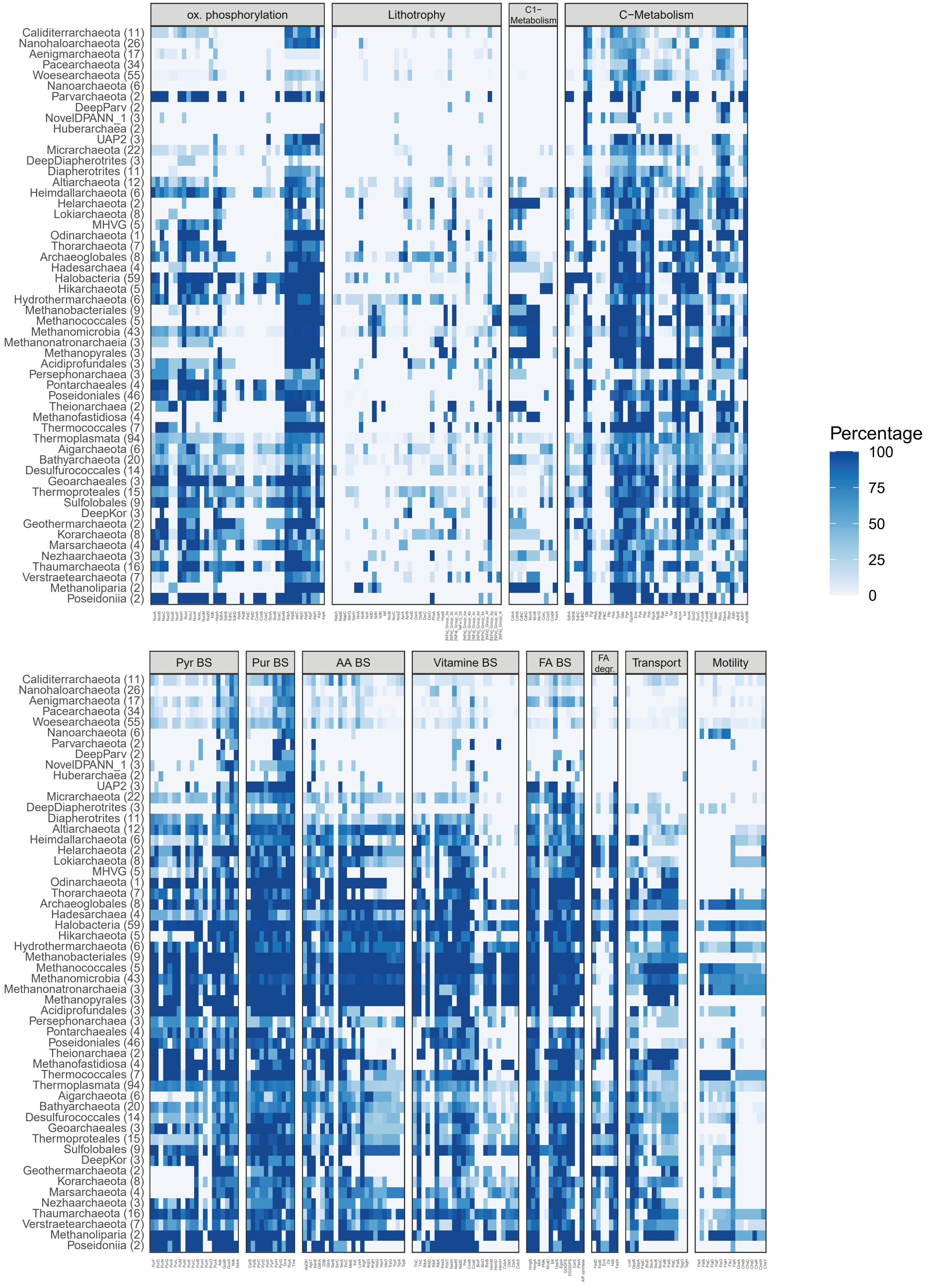


**Supplementary Figure 22 Presence of metabolic genes identified in archaeal genomes across the 651 genome dataset.** Heatmap shows presence/absence patterns of key metabolic genes across major archaeal groups included in the 651 genome dataset. Numbers in parentheses represent the number of genomes included in each major group. Supplementary Table 4 lists raw values used to produce the plot.

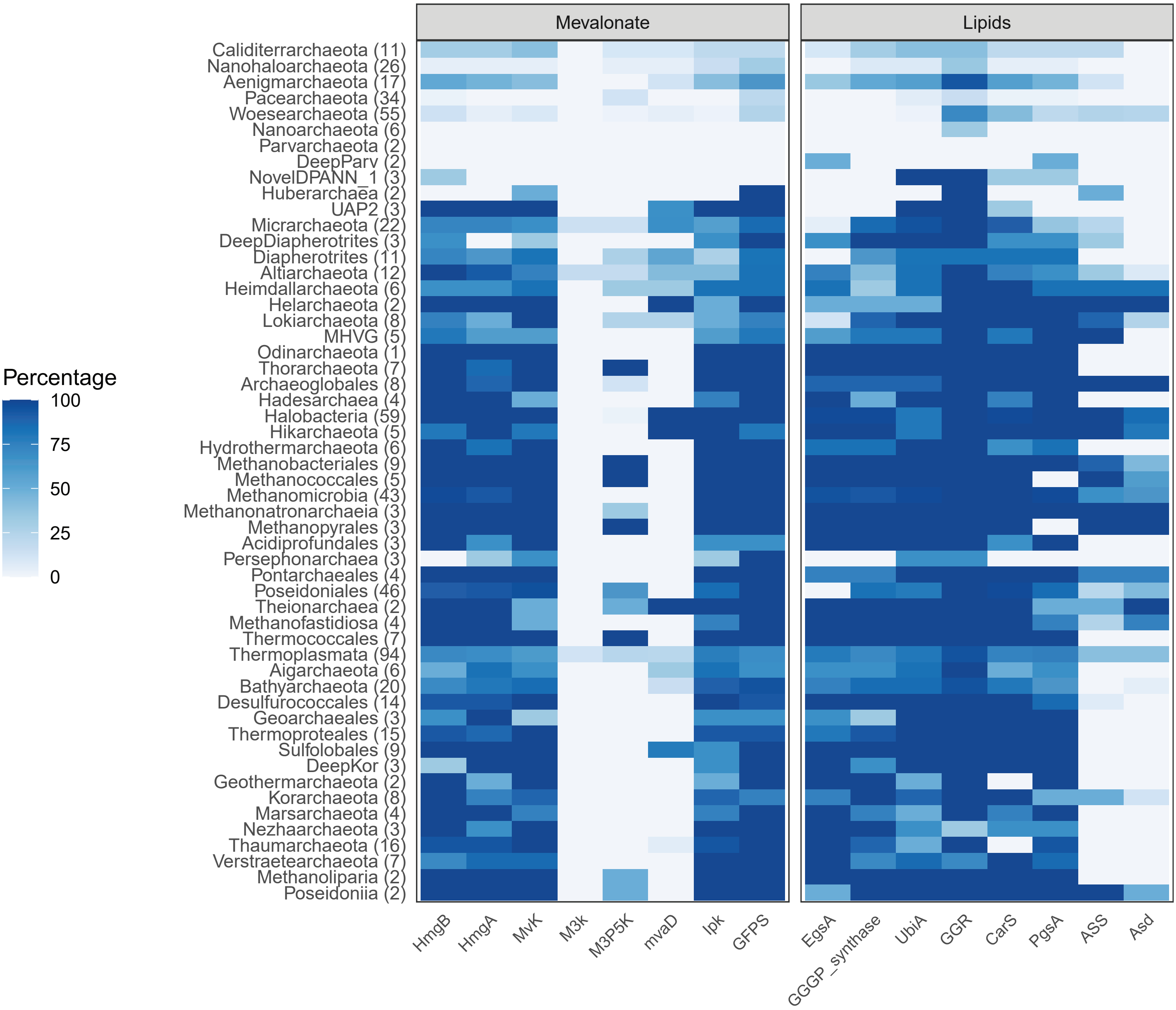


**Supplementary Figure 23 Presence of lipid biosynthesis genes identified in archaeal genomes across the 651 genome dataset.** Heatmap shows presence/absence patterns of key lipid biosynthesis genes across major archaeal groups included in the 651 genome dataset. Numbers in parentheses represent the number of genomes included in each major group. Supplementary Table 4 lists raw values used to produce the plot.

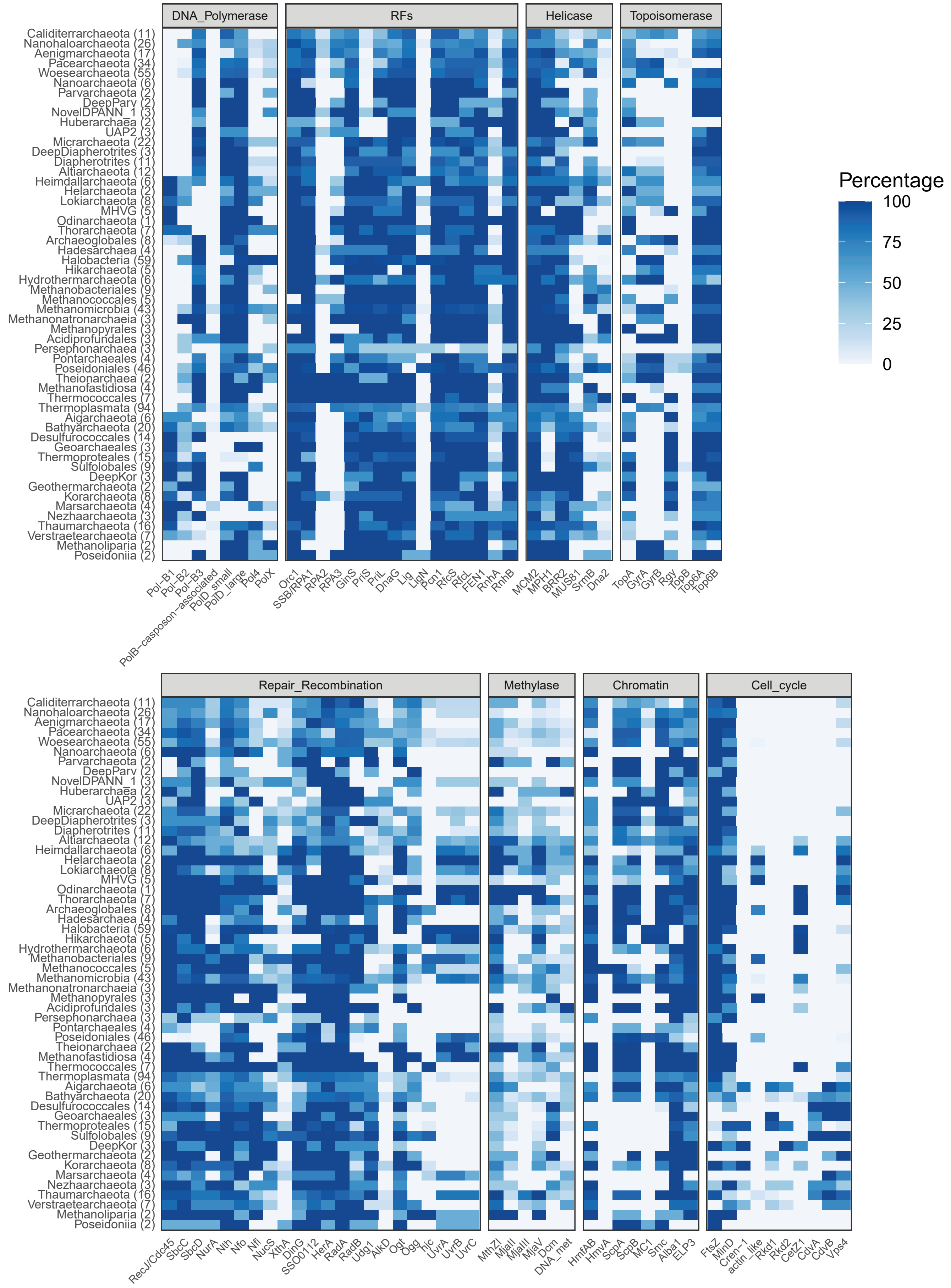


**Supplementary Figure 24 Presence of information processing and replication genes identified in archaeal genomes across the 651 genome dataset.** Heatmap shows presence/absence patterns of key information processing and replication genes across major archaeal groups included in the 651 genome dataset. Numbers in parentheses represent the number of genomes included in each major group. Supplementary Table 4 lists raw values used to produce the plot.

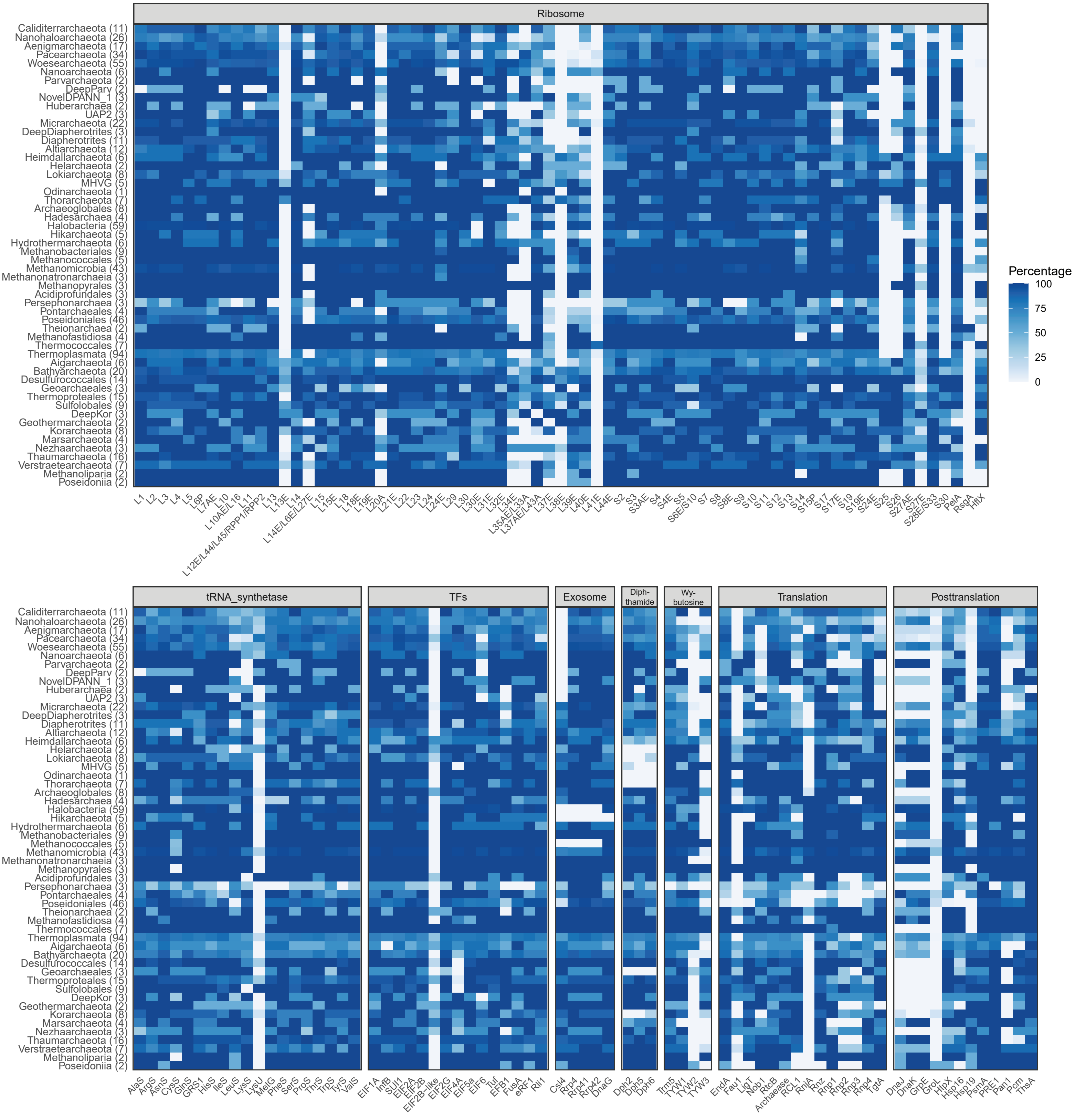


**Supplementary Figure 25 Presence of translation associated genes identified in archaeal genomes across the 651 genome dataset.** Heatmap shows presence/absence patterns of key translation associated genes across major archaeal groups included in the 651 genome dataset. Numbers in parentheses represent the number of genomes included in each major group. Supplementary Table 4 lists raw values used to produce the plot.

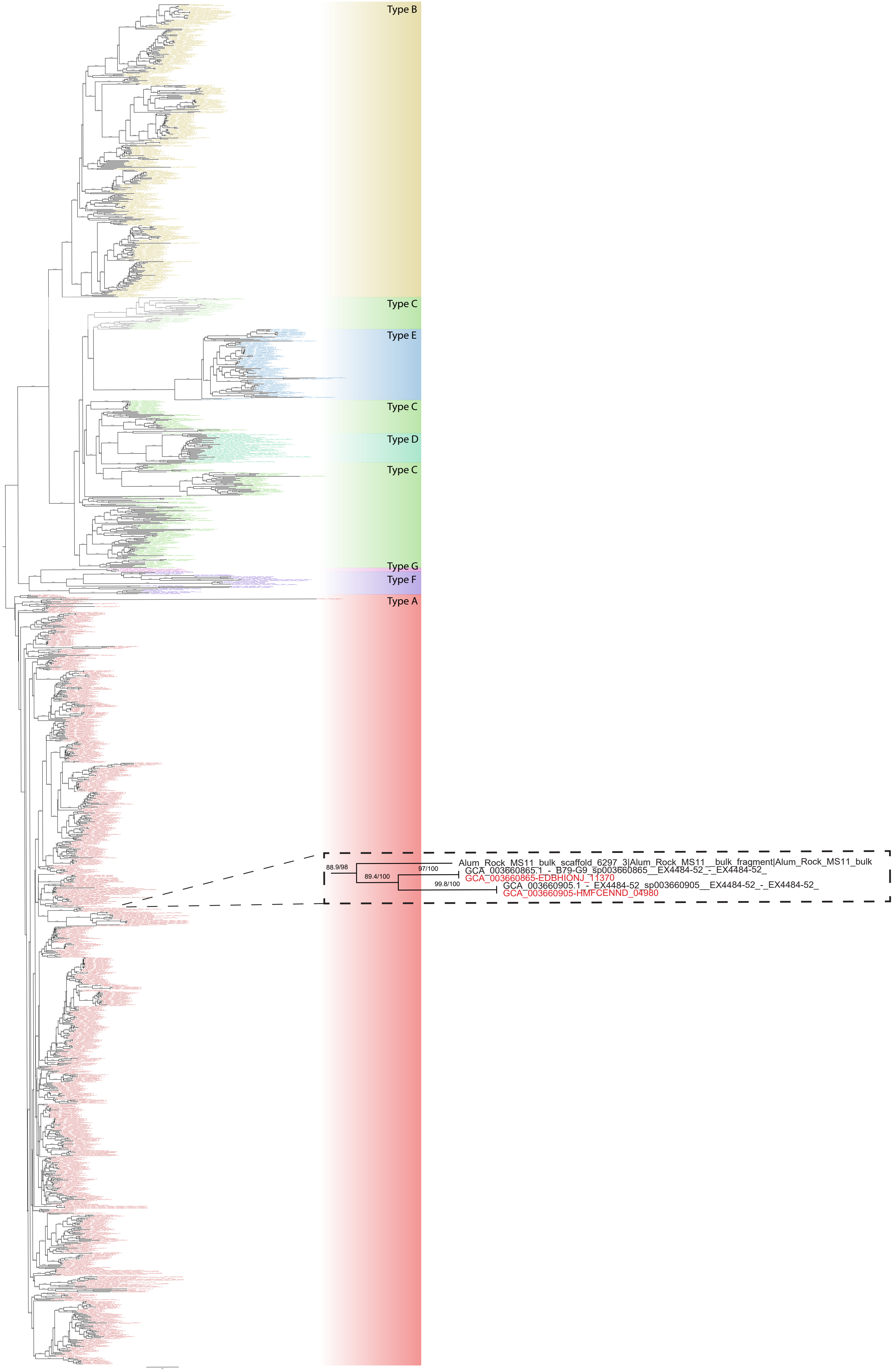


**Supplementary Figure 26 Phylogenetic placement of Caliditerrarchaeota FeFe Hydrogenase catalytic subunits.** Alignment was trimmed with BMGE (Alignment length = 141 aa). A ML phylogenetic tree was inferred using IQ-Tree with the LG+C20+F+R model and ultrafast bootstrap approximation (left) and SH-like approximate likelihood tests (right), each run with 1000 replicates. The tree has been artificially rooted at the midpoint. Inset highlights placement of Caliditerrarchaeota subunits within the Type A FeFe Hydrogenase cluster. Scale bar: Average number of substitutions per site.

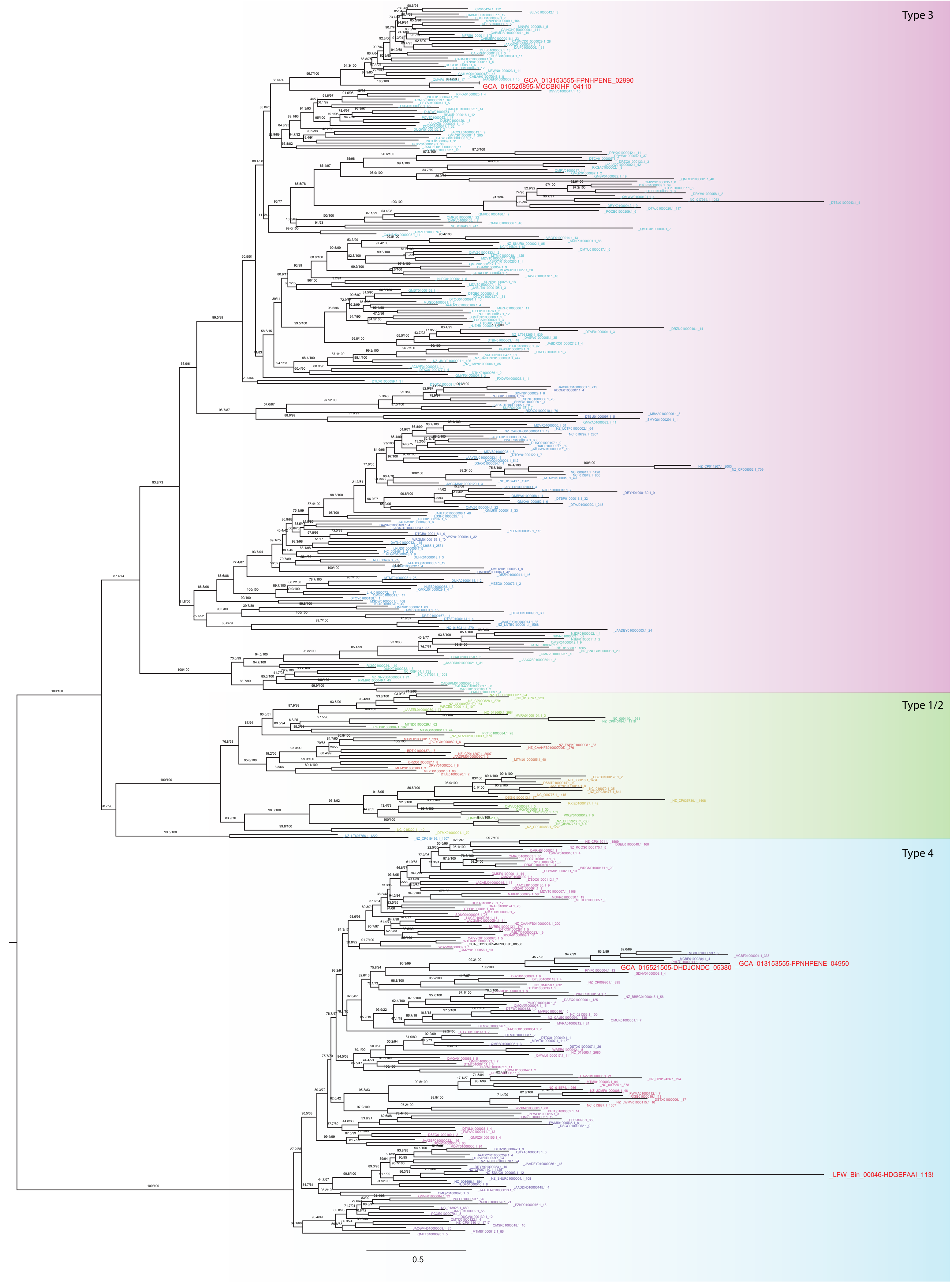


**Supplementary Figure 27 Phylogenetic placement of Calditerrarchaeota NiFe Hydrogenase catalytic subunits.** Alignment was trimmed with BMGE (Alignment length = 243 aa). A ML phylogenetic tree was inferred using IQ-Tree with the LG+C20+F+R model and ultrafast bootstrap approximation (left) and SH-like approximate likelihood tests (right), each run with 1000 replicates. The tree has been artificially rooted at the midpoint. Calditerrarchaeota genes are highlighted with red text. Scale bar: Average number of substitutions per site.

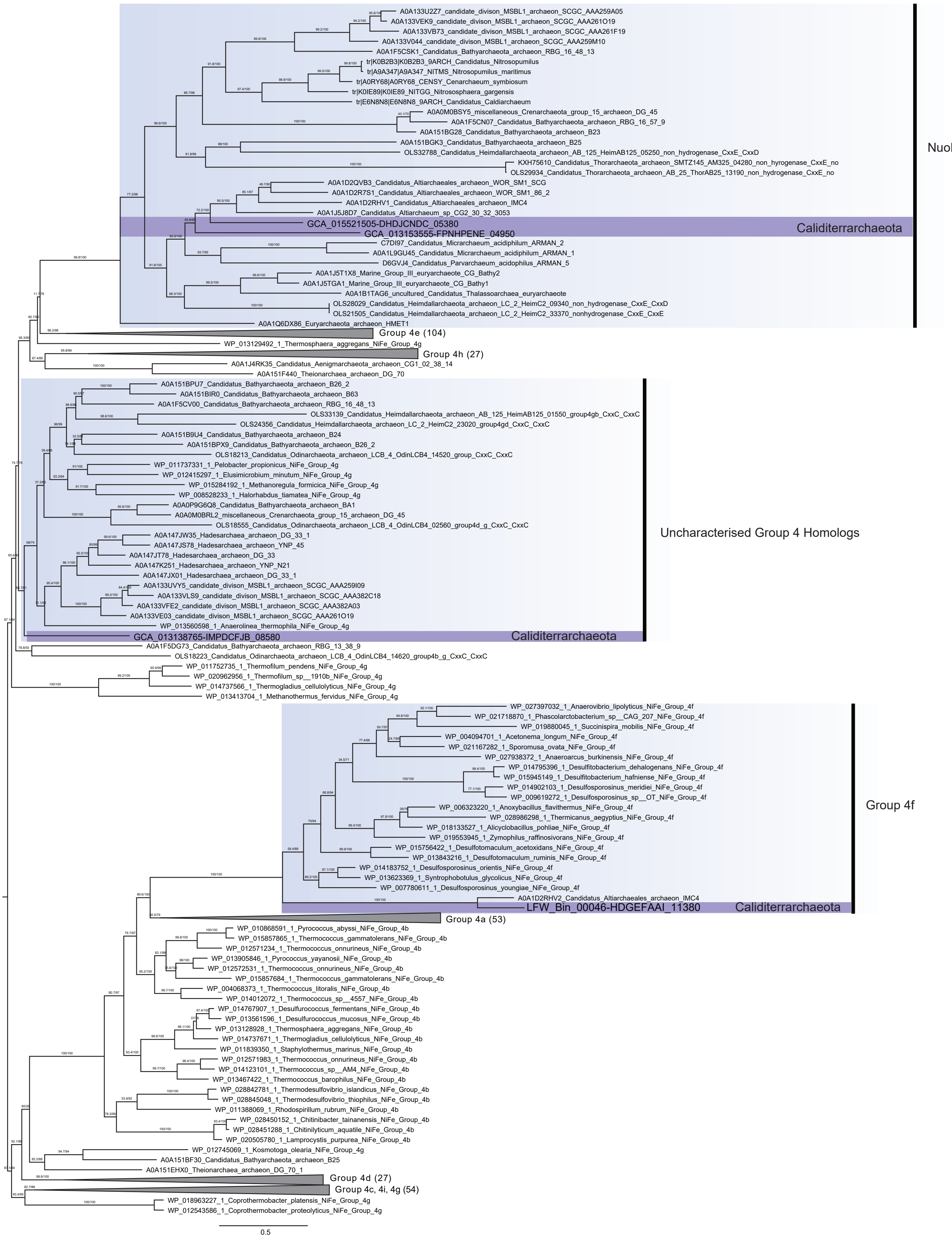


**Supplementary Figure 28 Phylogenetic placement of Calditerrarchaeota Group 4 NiFe Hydrogenase catalytic subunits.** Alignment was trimmed with BMGE (Alignment length = 324 aa). A ML phylogenetic tree was inferred using IQ-Tree with the LG+C20+F+R model and ultrafast bootstrap approximation (left) and SH-like approximate likelihood tests (right), each run with 1000 replicates. The tree has been artificially rooted at the midpoint. Calditerrarchaeota genes are highlighted. Scale bar: Average number of substitutions per site.

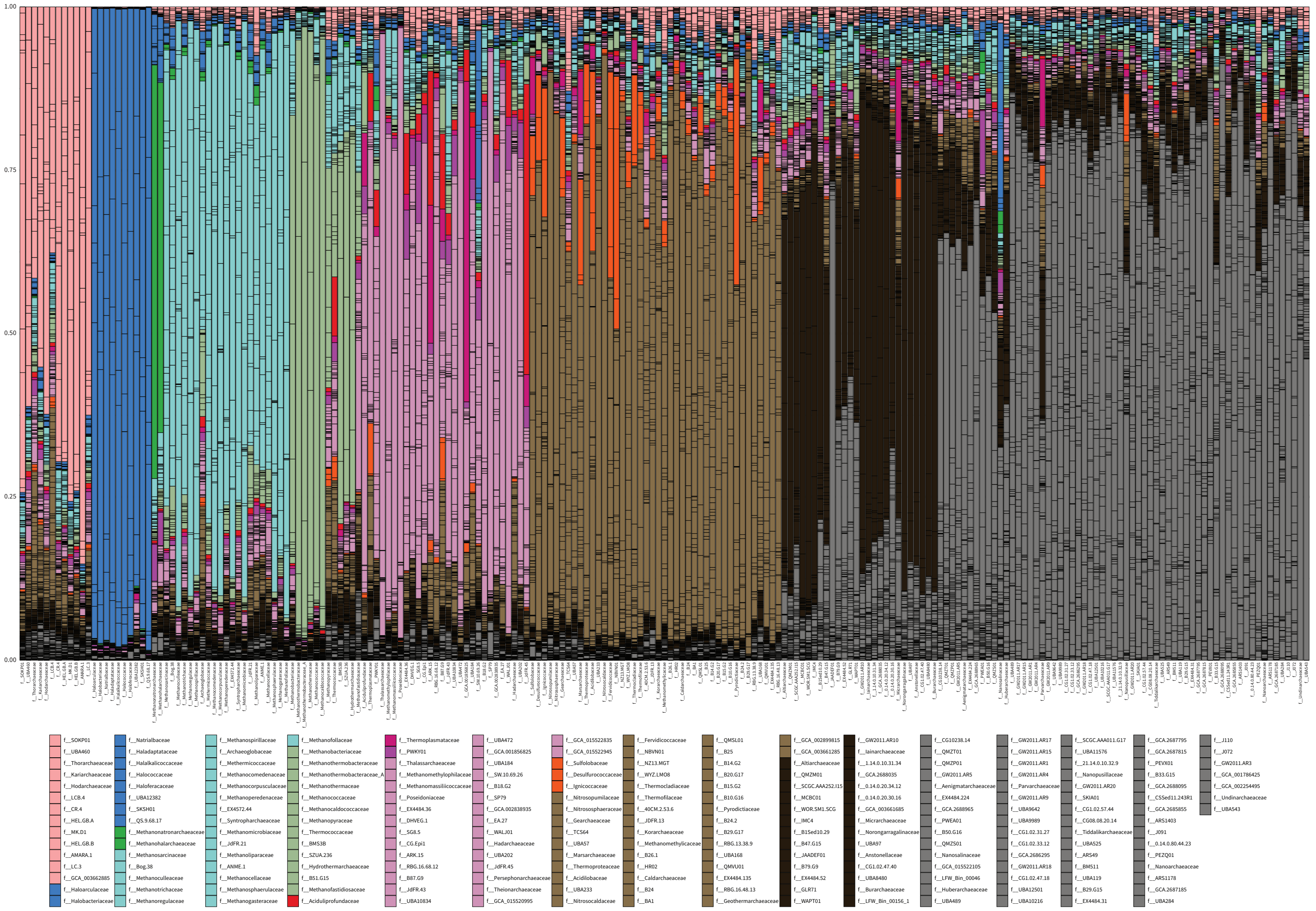


**Supplementary Figure 29 Rate of sisterhood for GTDB Family level lineages across 9,555 gene trees used for ALE analysis.** Relative sisterhood frequencies were calculated per gene and summed for each taxonomic group. Related lineages have been clustered together using colours and specific lineages that show higher rates of sisterhood with DPANN lineages highlighted with unique colours. The data used to generate this plot is available in Supplementary Table 11.
